# *Lactuca* super-pangenome reduces bias towards reference genes in lettuce research

**DOI:** 10.1101/2024.06.20.599299

**Authors:** Dirk-Jan M. van Workum, Sarah L. Mehrem, Basten L. Snoek, Marrit C. Alderkamp, Dmitry Lapin, Flip F.M. Mulder, Guido Van den Ackerveken, Dick de Ridder, M. Eric Schranz, Sandra Smit

## Abstract

Breeding of lettuce (*Lactuca sativa* L.), the most important leafy vegetable worldwide, for enhanced disease resistance and resilience relies on multiple wild relatives to provide the necessary genetic diversity. In this study, we constructed a super-pangenome based on four *Lactuca* species (representing the primary, secondary and tertiary gene pools) and comprising 474 accessions. We include 68 newly sequenced accessions to improve cultivar coverage and add important foundational breeding lines. With the super-pangenome we find substantial presence/absence variation (PAV) and copy-number variation (CNV). Functional enrichment analyses of core and variable genes show that transcriptional regulators are conserved whereas disease resistance genes are variable. PAV-genome-wide association studies (GWAS) and CNV-GWAS are largely congruent with single-nucleotide polymorphism (SNP)-GWAS. Importantly, they also identify several major novel quantitative trait loci (QTL) for resistance against *Bremia lactucae* in variable regions not present in the reference lettuce genome. The usability of the super-pangenome is demonstrated by identifying the likely origin of non-reference resistance loci from the wild relatives *Lactuca serriola, Lactuca saligna* and *Lactuca virosa*. The provided methodology and data provide a strong basis for research into PAVs, CNVs and other variation underlying important biological traits of lettuce and other crops.

## Background

Garden lettuce (*Lactuca sativa*) plays an important role in agriculture worldwide. In 2020, worldwide production of lettuce (including chicory) was about 27 billion tonnes, with a total value of about 20 billion US dollars (FAOSTAT, 2023). Because of its size and easy cultivation, lettuce is the perfect candidate for new farming techniques such as vertical farming, hydroponics and light-emitting diode (LED) illumination. In addition, lettuce is suitable for modern breeding techniques such as genome editing by Clustered Regularly Interspaced Short Palindromic Repeats (CRISPR)-Cas (Luo et al., 2021) and has many economically important close relatives within the Asteraceae such as endive, chicory, sunflower and chrysanthemum. Thus, lettuce has the potential to become a model organism for all Asteraceae.

For accurate, contemporary and inter-specific research into lettuce, a complete understanding of the lettuce genome is needed. Currently, a chromosome-level genome assembly is available for *L. sativa* (Reyes-Chin-Wo et al., 2017) as well as whole-genome sequencing (WGS) data for hundreds of *Lactuca* accessions (Wei et al., 2021). Lettuce genomic research has been centred around a single reference genome assembly for a long time, thereby likely introducing so-called “reference bias”. Reference bias is caused by ignoring differences in genomic composition and rearrangements between analysed accessions and the reference accession. Of particularly large impact are gene presence/absence variation (PAV), gene copy-number variation (CNV) and other large structural variations. Absence of regions in a reference assembly due to such variations can result in missing relevant information or incorrect genotyping. Moreover, it is known that variable genes (*i*.*e*. genes present in only a subset of accessions of a species) are enriched for functions important to agronomic traits such as disease resistance (Gao et al., 2019; Golicz et al., 2016; Y.-h. Li et al., 2014; Montenegro et al., 2017; Song et al., 2020; Zhao et al., 2020). This results in a strong need for expanding our vision beyond the single reference genome to get an integrated overview of genetic variation in both cultivated and wild *Lactuca* accessions. Pangenomics – the genetic study of closely related individuals first introduced by Tettelin et al. (2005) – does this by taking all available genomic variation among related individuals into account.

Similarly to other crops, the availability of large-scale sequencing efforts enables an integrated analysis of genetic variation among both cultivated and wild relatives of lettuce. This allows for the exploitation of genetic variation across species boundaries for the improvement of traits of interest. So far, only single-nucleotide polymorphisms (SNPs) and InDels with respect to *L. sativa* were analysed (Wei et al., 2021; Z. Zhang et al., 2023), whereas the wealth of WGS data al-for a more thorough investigation of PAV and CNV across species. Traditionally, wild relatives of a cultivated crop species have been classified into three gene pools based on their ability to cross with it (Harlan & de Wet, 1971). *Lactuca serriola*, being the evolutionary predecessor of *L. sativa*, belongs to the primary gene pool for lettuce breeding since there are no limitations for crossing with *L. sativa* (Lindqvist, 1960; L. Zhang et al., 2017). The secondary gene pool contains *Lactuca saligna*, which produces partly fertile offspring with *L. sativa*. Finally, the tertiary gene pool contains *Lactuca virosa*, which can only be crossed with *L. sativa* under specific circumstances (Lindqvist, 1960; Maisonneuve et al., 1995). For both *L. saligna* and *L. virosa*, near chromosome-level genome assemblies became available recently (Xiong, Berke, et al., 2023; Xiong, van Workum, et al., 2023). All of this data allows for the analysis of a lettuce super-pangenome, a pangenome of both cultivated and wild accessions from different species (Khan et al., 2020). Since its introduction, this concept has already been successfully applied to rice, lentil, maize, soy and tomato (Gui et al., 2022; Gutierrez-Gonzalez et al., 2022; N. Li et al., 2023; Shang et al., 2022; Zhuang et al., 2022).

Here, we describe the analysis of a 474-accession *Lactuca* super-pangenome comprising variation within the three gene pools available for lettuce breeding. As genes are the building blocks of the genome, we focus on the accurate identification of all genic PAVs and CNVs in accessions that have publicly available WGS data as well as newly generated WGS data for *L. sativa*. We built linear pangenomes per species for obtaining a full inventory of genes before identifying PAV and CNV. PAVs specifically allow for the biological interpretation of the core and variable genome of *Lactuca* species. Next, we show how PAV and CNV information can be used in genome-wide association studies (GWAS) to study genotype-phenotype associations, extending the traditionally used SNP-based analysis (Y. Sun et al., 2022; Zhao et al., 2020). Illustrating the vast amount of information in this super-pangenome resource, we describe a few key observations such as the make-up of the core and variable *Lactuca* genome, the use of PAVs to study the evolution of *Lactuca*, and non-reference resistance loci to *Bremia lactucae*.

## Results

### Constructing a *Lactuca* super-pangenome

Following Khan et al. (2020), a *Lactuca* super-pangenome across four species was constructed, combining variation found in 474 cultivated and wild accessions v (Supplementary Table S1, S2). This super-pangenome was built from four linear pangenomes, one for each of the four species relevant to lettuce breeding (*L. sativa, L. serriola, L. saligna, L. virosa*) (Figure 1). Per species, we combined the reference genome with non-reference sequence identified and assembled from low-coverage resequenced accessions of that species, and annotated the non-reference genes therein. No duplicated or contaminated sequences were added in the process (Supplementary Methods – Contamination filtering). For each species, we discovered between 10 and 18 Mbp non-reference sequence containing about 2,000 to 3,000 novel genes (Table 1). Finally, we quantified PAVs and CNVs from the alignment of WGS data per linear pangenome. A PAV is calculated as the fraction over the length of a predicted transcript that is covered by WGS reads. A CNV is calculated as the average depth over a transcript by WGS reads. This means that neither PAV nor CNV is a discrete value. However, some applications need discrete values for which we created a binary PAV (bPAV) matrix by applying a threshold.

**Table 1:** General statistics for all used reference genomes and the linear pangenomes created for these species. Note that the same reference genome was used for both *L. sativa* and *L. serriola*: as *L. serriola* lacks a high-quality reference genome and is closely related to *L. sativa*, we used the *L. sativa* reference genome. The total length is the total length of the assembly in basepairs. For all linear pangenomes, both total and the difference (Δ) compared to the genome of the same species are calculated.

| Assembly | Origin data | Total length ($\Delta$) | #sequences ($\Delta$) | #transcripts ($\Delta$) |
| --- | --- | --- | --- | --- |
| *L. sativa* genome | GCF_002870075.2 | 2,391,578,241 | 8,325 | 45,246 |
| " pangenome | " + 198 accessions | 2,401,543,966 (+9,965,725) | 14,458 (+6,133) | 48,689 (+3,443) |
| *L. serriola* genome | GCF_002870075.2 | 2,391,578,241 | 8,325 | 45,246 |
| " pangenome | " + 191 accessions | 2,408,948,407 (+17,370,166) | 18,928 (+10,603) | 48,505 (+3,259) |
| *L. saligna* genome | PRJEB56287 | 2,166,282,974 | 12 | 42,908 |
| " pangenome | " + 49 accessions | 2,184,875,879 (+18,592,905) | 10,509 (+10,497) | 46,484 (+3,576) |
| *L. virosa* genome | GCA_928212115.1 | 3,447,207,058 | 5,857 | 39,887 |
| " pangenome | " + 36 accessions | 3,459,562,374 (+12,355,316) | 13,510 (+7,653) | 41,743 (+1,856) |

**Figure 1:**
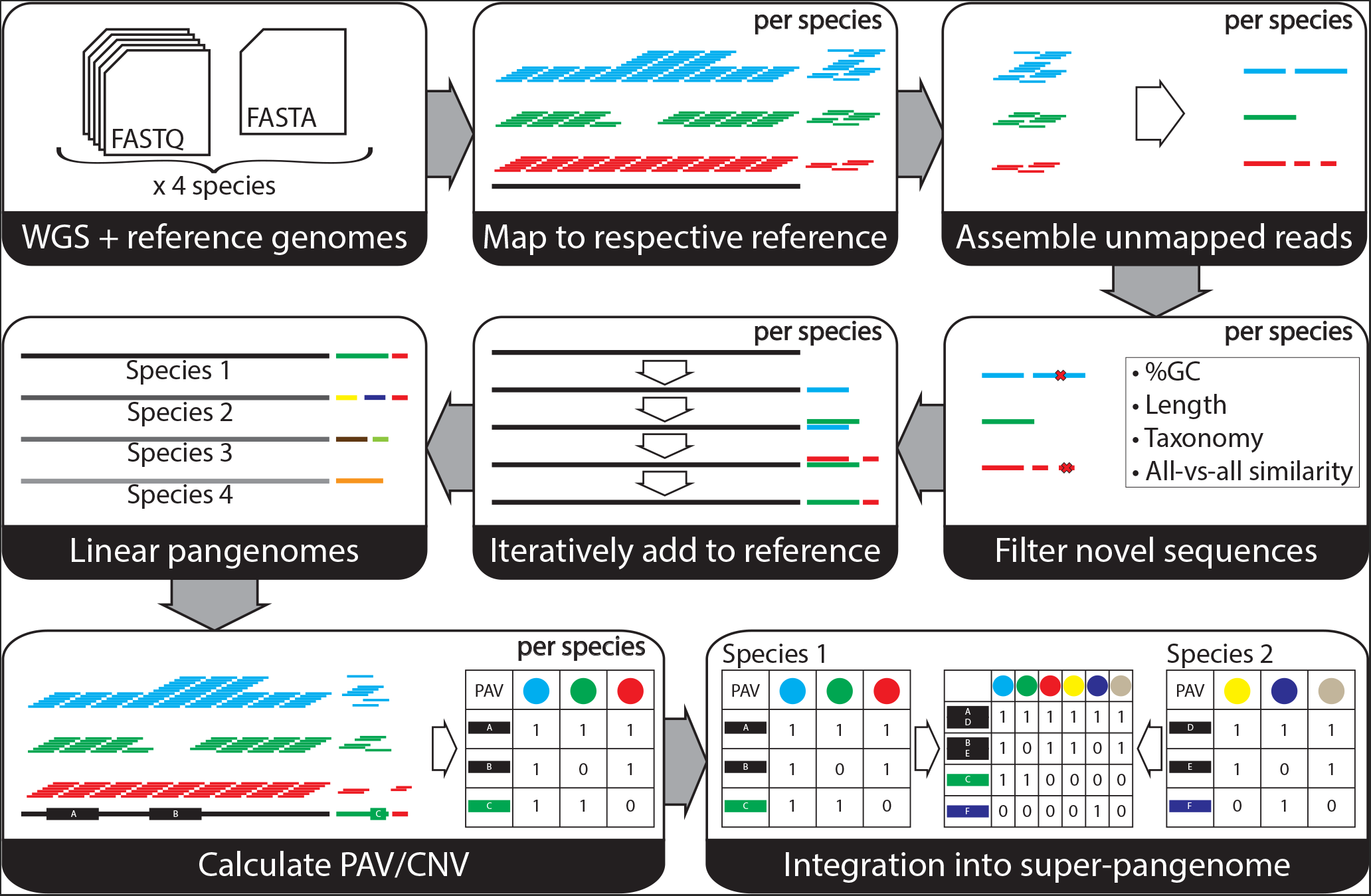
Overview of the process of going from WGS data for multiple species with their respective reference genomes to a super-pangenome in which all data is integrated across accessions and species. For each species separately, we mapped all available WGS sets per accession and extracted the unmapped reads. These unmapped reads were assembled and filtered for contamination, length and duplications. Then, we iteratively extended the reference genome to a linear pangenome with these novel sequences as to prevent any duplications. Mapping the WGS data per species against their respective linear pangenomes revealed presence/absence variation (PAV) and copy-number variation (CNV) values per transcript per accession. Finally, the linear pangenomes were integrated into a super-pangenome for all four species using PanTools.

These linear pangenomes were then combined in a super-pangenome graph database using PanTools (Jonkheer et al., 2022). Transcripts from all linear pangenomes (based on the proteins they encode) were grouped in homology groups, with settings optimised based on the distribution of Benchmarking Universal Single-Copy Orthologs (BUSCO) genes. Then, a super-pangenome bPAV matrix was created by combining the four linear pangenome bPAV tables based on homology (as illustrated in bottom right panel of Figure 1). The super-pangenome bPAV matrix thus indicates the number of genes present per homology group per accession. This resource enables gene set analyses, the comparison of PAVs and translation of biological information across species.

bPAV analysis shows that the variable part of the pangenome differs per species (Figure 2a). Based on the thresholds we set (see Supplementary Methods – PAV threshold for presence/absence), per species we found between 5,000 and 15,000 transcripts that show PAV (the so-called ‘variable genome’), which includes about 2,000 to 3,000 non-reference PAVs. Each core genome (per species) contains between 37,000 and 41,000 transcripts. Although the total pangenome size for *L. sativa* and *L. serriola* is comparable, *L. serriola* has a smaller core pangenome size and thus exhibits more variation between accessions. We hypothesise this is because *L. sativa* has potentially gone through a genetic bottleneck, which has removed (presence/absence) variation (Wei et al., 2021). Specifically compared to *L. saligna*, which has about the same sample size as *L. virosa, L. virosa* exhibits very little PAV. This corresponds to the geographical sampling spread of these species: the *L. virosa* accessions sampled are all located in Western Europe, whereas *L. serriola* and *L. saligna* accessions were sampled in both Europe and Asia (Wei et al., 2021).

**Figure 2:**
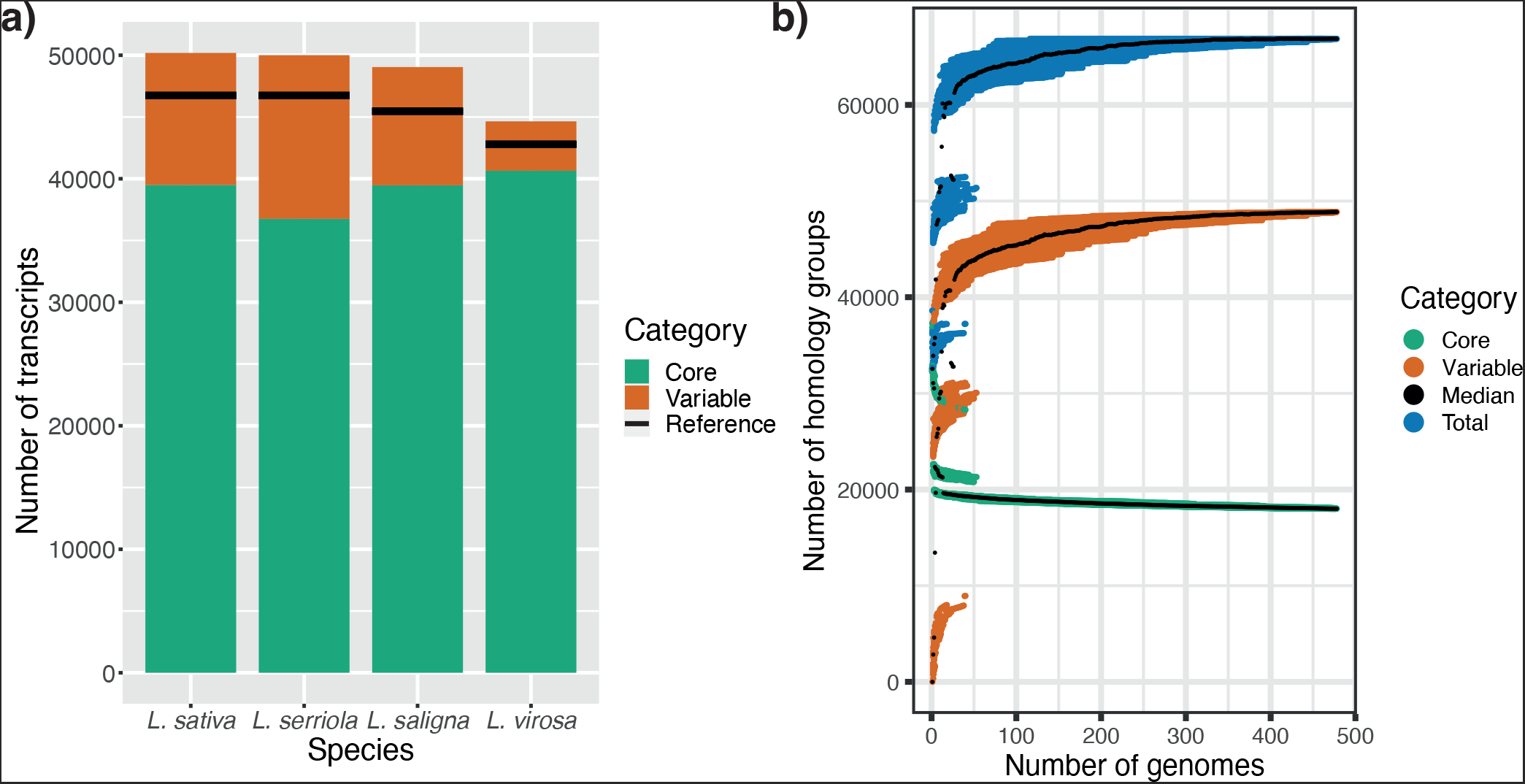
**a**) Pangenome size for each of the *L. sativa, L. serriola, L. saligna* and *L. virosa* linear pangenomes. **b**) Growth curve of the super-pangenome constructed from combining the four linear pangenomes. This growth curve is created by calculating core, variable and total homology group size for random iterations over the accessions in the super-pangenome. This process was repeated 100 times.

Across the four species in the *Lactuca* superpangenome, bPAV analysis reveals a clear distinction between the three gene pools and shows that about 50% of genes are shared across these pools (Figure 2b, Supplementary Data 7). In the super-pangenome growth curve, two clear jumps can be seen that occur when crossing phylogenetic boundaries. This corresponds to the three distinct gene pools present: (1) *L. sativa* and *L. serriola*, (2) *L. saligna* and (3) *L. virosa*. Also, the number of core homology groups quickly converges to about 18,000, indicating that the current set of accessions provides a good view on the core genes that make up *Lactuca*. The jumps in the variable genome size, on the other hand, are larger than for the core genome. Nevertheless, the Heaps’ Law alpha value (1.09) is higher than 1, indicating a closed pangenome (Tettelin et al., 2008).

### PAVs enable transfer of biological knowledge across species

The super-pangenome facilitates integration of genomic information across species. This allows for high-level analyses such as those of gene sets, but also provides the possibility to approximate the phylogeny of all accessions without the need for variant calling and to perform detailed analyses such as the translation of gene knowledge between species.

The 18,009 core homology groups contain about half of the genes in each accession. Interestingly, this is not a large deviation from what we previously reported on a three genome assembly comparison between *L. sativa, L. saligna* and *L. virosa* (Xiong, van Workum, et al., 2023). A functional enrichment of the core genes mainly identified transcription factor (TF)-related domains (Supplementary Figure S1). This shows that TFs, often master regulators of essential processes, play a role in defining the *Lactuca* genus.

The variable part of genomes of other crops are often enriched for disease resistance genes. These genes are known to be highly variable, both in terms of SNPs and InDels, and in terms of larger variation such as PAVs (Tamborski & Krasileva, 2020). This is relevant to plant breeding, since crops need to be protected against ever-evolving pathogens. The available variation among resistance genes provides breeders material to combat the evolution of pathogens. We found the variable genome of *Lactuca* to be not only significantly enriched for resistance genes but also for transposon-related genes (Supplementary Figure S2, Supplementary Data 9). Transposons are known to be both highly variable and a source of phenotypic variation (Cai et al., 2022; Golicz et al., 2016). Some major traits in plants are shown to be caused by transposons, *e*.*g*. the insertion of a transposon in *LsKN1* (*L. sativa knotted 1*) is responsible for head formation in lettuce (Yu et al., 2020).

As SNPs are genomic markers of evolutionary history, they can be used for a good approximation of the phylogeny of a species. Therefore, we reason that PAV should contain similar information. PAVs – as opposed to SNPs – do not involve variant calling and take non-reference genomic content into account. We built a neighbour-joining tree for all accessions in the superpangenome based on bPAV information (Figure 3). The *L. serriola* clade in the tree is very similar to the SNP-based tree by Wei et al. (2021), which was shown to reflect geographic origin. Both *L. saligna* and *L. virosa* clades also closely follow geographic distribution. We were able to distinguish most types (also called: sub-groups) of *L. sativa* accessions. Especially the main breeding types Butterhead, Oilseed and Crisp lettuce clearly cluster together based on PAVs.

**Figure 3:**
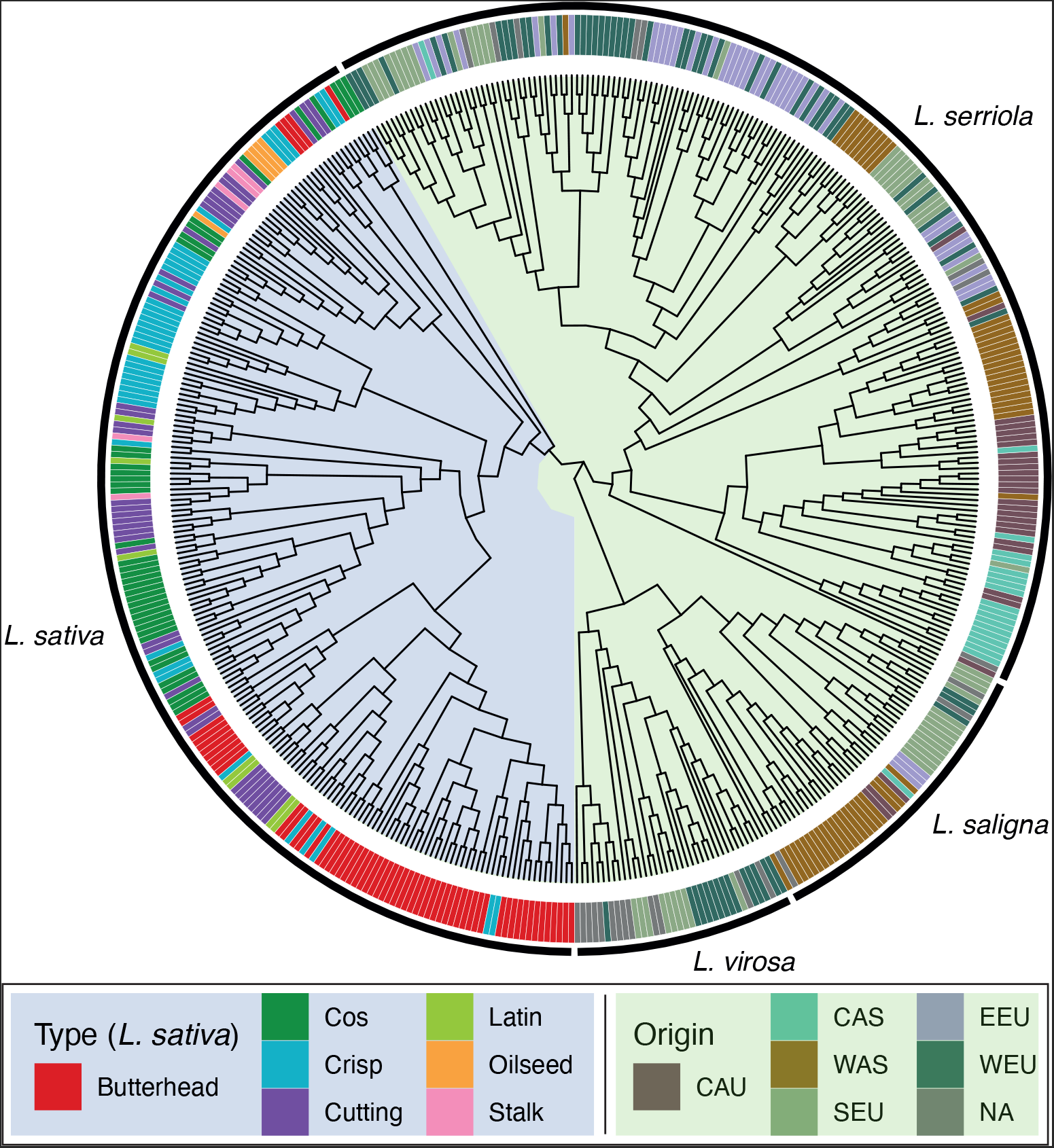
The super-pangenome bPAV matrix was used to construct a neighbour-joining tree of 474 *Lactuca* accessions. To indicate relatedness of accessions, type (also knows as ‘subgroup’) names are used for *L. sativa* and geographical origin for the wild lettuce relatives. The classification of origin as CAU (Caucasus), CAS (Central Asia), WAS (Western Asia), SEU (Southern Europe), EEU (Eastern Europe) and WEU (Western Europe) was taken from Wei et al. (2021).

### PAV-GWAS and CNV-GWAS are complementary to SNP-GWAS

To investigate where PAVs and CNVs are associated to phenotypes, we used the PAV and CNV calls as genotypes for GWAS. First, we performed a validation with the locus that is responsible for variation in seed colour. This locus has previously been identified with SNP-GWAS and a SNP in the the *LsTT2* gene has subsequently been confirmed to be the cause of the black/white phenotype (Kwon et al., 2013; Mehrem et al., 2024; Seki et al., 2024; X. Zhang et al., 2023). With PAV-GWAS we indeed find the same locus for seed colour as traditional SNP-GWAS. As expected, *LsTT2* itself shows no PAV (the cause is a SNP) but neighbouring genes that do enable us to find the locus (Figure 4a). Second, Su et al. (2020) found a CNV of the *RLL2B* gene to cause a difference in anthocyanin content in *L. sativa* plants, with a tandem duplication in red accessions but only one copy in green accessions. This duplication locus was previously found with traditional SNP-GWAS and is confirmed in our CNV data (Supplementary Figure S3) (Wei et al., 2021). Specifically, when running CNV-GWAS with the anthocyanin phenotypes available through the database from the Centre for Genetic Resources, the Netherlands (CGN), we found two neighbouring CNVs of *RLL2B* on chromosome 5 to be significant (both correlate with the phenotype and *RLL2B*; thus identifying the correct locus (Figure 4b)).

**Figure 4:**
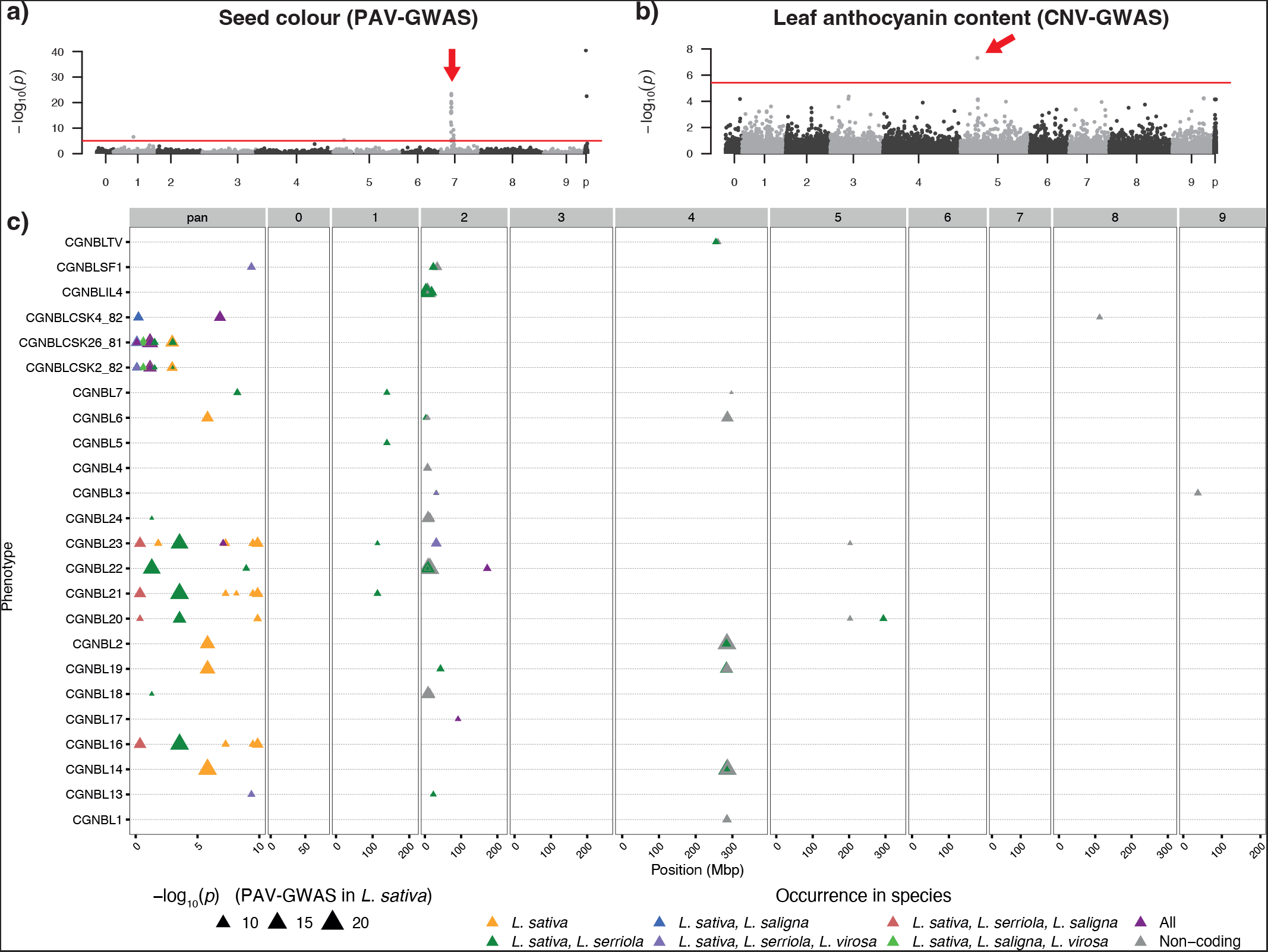
Manhattan plots for PAV-GWAS and CNV-GWAS show they are complementary to SNP-GWAS. For each PAV, only the longest isoform per gene is visualised. In **a**), we show a clear association between seed colour of *L. sativa* and PAV genotypes that was found previously using SNP-GWAS (X. Zhang et al., 2023). **b**) shows that CNV-GWAS can be used for the discovery of the locus responsible for the leaf anthocyanin phenotype (Su et al., 2020). In both **a**) and **b**), the red line indicates the significance threshold according to Bonferroni multiple testing correction (™ log_10_(*p*) > *threshold*). Red arrows indicate the peaks mentioned in the main text. **c**) PAVs in non-reference genes are highly associated with resistance to specific races of lettuce downy mildew *B. lactucae*. The phenotype is the severity score as observed by CGN. Only significant PAVs (after Bonferroni multiple testing correction: ™ log_10_(*p*) > 5.3) are shown, with marker colour indicating occurrence in other species of the super-pangenome and size corresponding to *p*-value. As non-coding genes cannot be used for homology grouping, they are coloured grey.

Next, we applied PAV-GWAS to a set of publicly available phenotypes of susceptibility scores of *L. sativa* accessions to *B. lactucae* races from CGN. Resistance to pathogens, such as the downy mildew *B. lactucae*, is mostly conveyed by nucleotide-binding leucine-rich repeat, and in more exceptional cases by receptor-like kinase and receptor-like protein families (Parra et al., 2016). In lettuce, these genes are commonly found in major resistance clusters that are known to display PAV (Christopoulou et al., 2015; Shen et al., 2006). Confirming this, we found loci in PAV-GWAS of *B. lactucae* infection phenotypes mostly located in the known major resistance cluster of chromosome 2 (Figure 4c). This is in agreement with what was found previously with SNP-GWAS (Wei et al., 2021) (*cfr*. their Supplementary Figure 11). Furthermore, 36 of all 137 transcripts whose PAV is significantly associated with *B. lactucae* phenotypes do not appear in the reference genome, illustrating the potential impact of reference bias and underlining the importance of a pangenome (Supplementary Table S3). These PAV-containing loci provide starting points for further investigation of these regions.

One of the 36 significant, non-reference transcripts to *B. lactucae* was classified as a disease resistance protein: *g3691*.*t1*. The encoded protein has a TIR-NBS-LRR domain (PTHR11017:SF386) according to PANTHER. Since the presence of *g3691*.*t1* is associated with resistance, it could be the causal gene for resistance against *B. lactucae* BL23. From the super-pangenome, we know that this transcript is present in all four species, but in *L. sativa* and *L. serriola* it is absent from their reference genome. The homologues of this gene in *L. saligna* and *L. virosa* are both located at the start of chromosome 2, indicating that this gene is likely part of the major resistance cluster MRC2, which complicates the use of this finding in lettuce breeding.

The non-reference gene whose transcript was significantly associated to most *B. lactucae* phenotypes was *g767*. The PAV pattern of this transcript is significantly associated to 5 of the 24 *B. lactucae* phenotypes collected from the CGN database: CGNBL16, CGNBL18, CGNBL20, CGNBL21, CGNBL23 (Figure 4c). This transcript is present in 19 of the 198 *L. sativa* accessions and its presence is always negatively correlated to *B. lactucae* susceptibility, making it a potential resistance locus (Supplementary Table S3). Using correlation with SNPs, we determined this gene likely to be located within the major resistance cluster of chromosome 2 as well. No functional domains could be annotated for the encoded protein by InterProScan. However, in the *L. serriola* pangenome we found a homologue of this gene that is present in 13 of the 191 *L. serriola* accessions, possibly the origin of this locus.

Similarly to SNP-GWAS, GWAS based on CNVs/PAVs is able to identify a locus significantly associated to a phenotype if the causal gene is either a CNV/PAV itself or in linkage disequilibrium (LD) with a CNV/PAV. This makes CNV-GWAS and PAV-GWAS complementary to SNP-GWAS, as CNVs do not necessarily contain SNPs and SNPs are impossible to call for absent regions. Finally, the super-pangenome (including cultivated and wild lettuce) enables us to trace back the potential origin of genes.

## Discussion

We described the analysis of a 474-accession *Lactuca* super-pangenome, spanning four species. We demonstrated the added value of a pangenome for GWAS based on gene presence/absence and copy-number variations over reference-based SNP-GWAS. Also, the super-pangenome enabled us to functionally characterise (species-specific) core and variable sets of genes for *Lactuca*. It thereby allowed for reasoning across and comparison between wild and cultivated *Lactuca* species. Below, we discuss a number of general findings.

### Quality matters for a reliable pangenome

The quality and completeness of a pangenome critically depend on its underlying data. Thus, from the publicly available genome assemblies we selected those with the highest quality as a reference for mapping and PAV determination, even though this resulted in using the *L. sativa* genome for *L. serriola* accessions (Supplementary Methods – Data selection). At the time of analysis, we used the only available genome assembly, GCF_002870075.2 (*L. sativa* var. Salinas v8). The most recent version (v11: GCF_002870075.4) is a novel genome assembly based on HiFi sequencing instead of Illumina and consequently contains almost no gaps. However, in terms of genic content v11 is almost as complete as v8 (Supplementary Figure S4).

The linear pangenomes are constructed from shortread data, which has drawbacks for the quality of the resulting pangenome. The assembly of all reads that did not map to the high-quality reference genome contains genomic material of other species than only lettuce. Even though this contamination should not be present in a high-quality reference genome, it will be assembled as novel content in the linear pangenome. For separating *Lactuca* from non-*Lactuca* content, several metrics were used (see Supplementary Methods – Contamination filtering). Nevertheless, a number of accession-specific PAVs for some accessions in the final PAV analysis are potentially due to unknown contamination. Given short-read data only, we consider it impossible to design a general-purpose pipeline to construct linear pangenomes that are complete, non-redundant and fully free of contamination.

To unequivocally assess homology relationships, ideally all gene sequences in an accession should be available. This requires complete genome assemblies for each accession, which is difficult given only short-read data. However, with the current surge in HiFi sequencing, we expect more of such complete assemblies to become available for *Lactuca* in the near future.

### The synergy between PAVs/CNVs and SNPs for GWAS

Even though a genotype may be significantly associated to a specific phenotype by GWAS, this does not imply a causal relationship. In SNP-based GWAS, LD causes SNPs genetically linked to the actual causal genotype to be significantly associated to a phenotype as well. The same is true for PAV- and CNV-GWAS, where LD is clearly visible for *e*.*g*. seed colour (Figure 4a). Other studies where PAVs were used for GWAS showed the presence of LD as well (Liu et al., 2022; Song et al., 2020). We found that PAV- and CNV-GWAS peaks overlap loci previously identified by SNP-GWAS in Mehrem et al. (2024) and Wei et al. (2021). The causal genotype may be the significant PAV or CNV (as *e*.*g*. shown by Song et al. (2020)), but confirmation still requires experimental validation.

An important novelty in this study is the use of continuous genotype values for PAVs and CNVs. In traditional SNP-GWAS, but also in other PAV-GWAS studies (Liu et al., 2022; Song et al., 2020; Y. Sun et al., 2022; Zhao et al., 2020), presence/absence and the number of copies of a transcript are quantified as integer values. Thresholds are used to distinguish presence from absence, *e*.*g*.: The software SGSGeneLoss used a threshold of at least 5% coverage and a minimum depth of 1 for all exons of a transcript (Golicz et al., 2015); the rice pangenome reported in C. Sun et al. (2017) used 95% coverage for the coding region and 85% coverage for the genic region as threshold; the tomato pangenome by Gao et al. (2019) used 20% coverage and a minimum depth of 2 for all exons of a transcript; and the pipeline Panoramic used 75% coverage or 50% coverage and a minimum depth of 3 (depending on quality) for the entire genic region (Glick & Mayrose, 2021). These choices unnecessarily simplify the underlying data and involve a sometimes rather arbitrary selection of thresholds. Since GWAS is based on a linear model and does not require independent variables to be integer, we made use of the continuous PAV and CNV values as computed from the underlying short-read alignments (see Supplementary Methods – PAV and CNV calculation) to avoid the use of an arbitrary threshold.

### Pangenomics is the future for lettuce breeding

A pangenomic approach makes it possible to identify novel genes and find associations between phenotypes and genotypes that are not necessarily present in the reference genome. The power of pangenomics lies in the possibility of combining all types of variation – SNPs, PAV, CNV and larger structural variations –, use it to identify novel breeding targets and interpret them across accessions or even species. The linear pangenomes and the analyses introduced here provide an interesting first look into the *Lactuca* super-pangenome, although ideally the need for the intermediate linear pangenomes will disappear with improved methodology.

To bring pangenomics to full application in plant breeding, however, further investigations are necessary. Firstly, expanding pangenomics to include other wild relatives of lettuce (such as *Lactuca dregeana, Lactuca aculeata* and *Lactuca altaica*) will provide a more comprehensive understanding of its genetic diversity. Additionally, generating and including additional genome assemblies will enable researchers to explore intergenic regions, inversions and translocations in detail. These will allow us to help confirm and physically position the here identified non-reference sequences. With the availability of more accurate and complete genome assemblies, we anticipate a clearer picture of (highly) variable regions within the *Lactuca* genome and their biological implications. Such insights can inform targeted breeding strategies and contribute to the development of improved lettuce varieties.

## Methods

### Data selection

We selected a total of 474 lettuce accessions (belonging to *L. sativa, L. serriola, L. saligna* and *L. virosa*) for analysis (Supplementary Methods – Data selection). Both publicly available and in-house WGS data were used as well as nuclear, mitochondrial and chloroplast genomes for each species (Supplementary Table S1). The accession numbers of the 406 WGS data sets used from Wei et al. (2021) can be found in Supplementary Table S2 (column ‘BGI sequencing’). Importantly, the *L. sativa* reference genome was used as reference genome for *L. serriola*. Only for *L. virosa*, a different accession was used for its mitochondrial genome, since none was available for the reference accession and only little sequence variation exists within lettuce mitochondrial genomes (Fertet et al., 2021).

### Sequencing

We grew and sequenced 68 accessions that are not part of the WGS data published by Wei et al. (2021) to increase the resolution of our dataset (Supplementary Table S2; column ‘LK sequencing’). Seeds were obtained from CGN and multiplied by single-seed descent, except for *L. saligna* 275-5 (PV15242) which was multiplied by regular propagation. Plants were grown on rockwool cubes under long day conditions (21°C/19°C, 16h light, ∼ 100 *µ*mol/m^2^/sec). The plants received 1x Hyponex on the day of sowing, 0.5x Hyponex on day 7, and tap water on the other days. Samples of the first and second true leaves were collected at 17 days in the form of two 10 mm leaf discs for two plants per accession. These were snap-frozen in liquid nitrogen and stored at -80°C. DNA extraction was performed with MagMAX™ plant DNA isolation kit using a Kingfisher robot. Library preparation was performed using the Truseq DNA nano protocol and paired-end sequencing (2×150 bp) was performed on the Illumina NovaSeq6000 machine with an S4 flowcell at Utrecht Sequencing Facility (USEQ).

### Linear pangenome construction

A custom pipeline was built based on the “iterative map and build” approach (Golicz et al., 2016), partly inspired by the methods described by Glick and Mayrose (2021) and Hu et al. (2017) (Figure 1, Supplementary Methods – Pipeline construction strategy). After trimming the input paired-end Illumina reads with Trimmomatic v0.39 (Bolger et al., 2014), each dataset was mapped to its closest reference genome with bwa-mem2 v2.1 (Vasimuddin et al., 2019). Using samtools v1.11 (H. Li et al., 2009), the alignments were sorted, merged and indexed for extracting all unmapped reads per accession. These unmapped reads were assembled per accession with MEGAHIT v1.2.9 (D. Li et al., 2015); in the resulting assemblies, contigs with a length below 1kbp, a GC percentage higher than 50% or a kraken2 v2.1.2 (Wood et al., 2019) classification of Bacteria, Arthropoda and Amoebozoa were removed. This mapping/assembly stage was performed in parallel to speed up processing of hundreds of samples.

Next, a linear pangenome was built by iteratively adding the newly assembled sequences to the reference genome. Whether a new sequence was already part of the pangenome was determined by performing a minimap2 v2.22 (parameters ‘-cxasm5 --secondary=no’) (H. Li, 2018) alignment of the new genomic content per accession to the previous pangenome build. Whenever an alignment was found, only the longer sequence was retained. Then, all new sequences in the pangenome were clustered with CD-HIT-EST v4.8.1 (W. Li & Godzik, 2006) at 90% similarity to further remove possible duplicates. Finally, blobtools v1.1.1 was run per set of novel sequences in a linear pangenome according to standard instructions and based on its output we only kept contigs with at least 10X coverage and either “no-hit” or “Steptophyta” taxonomy.

Before starting annotation, we created a repeat database per linear pangenome with RepeatModeler and used it for masking the novel sequences with Repeat-Masker (both tools run in the TETools v1.5 container) (Tarailo-Graovac & Chen, 2009). With PanTools v3.3.4, we retrieved the core set of proteins shared between the *L. sativa, L. saligna* and *L. virosa* reference genomes (Jonkheer et al., 2022; Sheikhizadeh Anari et al., 2018). With only these core genes, we trained a new AUGUSTUS v3.4.0 model for *Lactuca*, which we applied to all novel sequences in the pangenome for gene prediction (Stanke & Waack, 2003). Finally, we searched all newly predicted proteins in the nr database with mmseqs2 v13.45111 and only kept contigs that had at least one hit to a species with taxonomy “unknown” or “Eukaryota” (Steinegger & Söding, 2017).

The full pipeline was implemented in Snakemake (Köster & Rahmann, 2012) and is available via: https://github.com/LettuceKnow/linear_pangenome_building.

### Presence/absence and copy-number variation analysis

For calling PAV and CNV of genes, we mapped all WGS data to the linear pangenome of the same species (*i*.*e*. closest genome) with bwa-mem2. Horizontal coverage (breadth; the fraction of the length of a transcript covered by at least one read) was calculated as a proxy for PAVs. Vertical coverage (depth; the average sequencing depth over a transcript divided by the median value of all average sequencing depths) was calculated as a proxy for CNVs. We calculated these metrics with bedtools ‘coverage’ (v2.30.0) (Quinlan & Hall, 2010), only considering exon locations. Metrics on exons were combined into metrics for transcripts based on their “Parent” attribute according to GFF3 information, thus ignoring coverage information for intronic regions. Organellar genes were excluded from these analyses.

For some downstream analyses, we could not make use of continuous PAV values. In these cases, we “binarised” the PAV values by calling genes present when PAV > 0.8 and absent otherwise (see Supplementary Methods – PAV threshold for presence/absence). This binary PAV matrix we call bPAV.

### Functional analysis

For each linear pangenome, only the longest isoform per gene was functionally annotated. Extracting the longest isoforms and translating these to protein sequences was done with AGAT v0.5.0 (Dainat et al., 2021). InterProScan v5.53-87.0 (Jones et al., 2014) was used for functional annotation. Next to searching for associated InterPro domains, GO terms and pathways, we searched for TIGRFAM, SUPERFAMILY, PANTHER, Gene3D, Coils, Pfam and MobiDBLite annotations.

### Super-pangenome construction

The four linear species panproteomes (using protein sequences only) were integrated into one genic superpangenome using PanTools (Jonkheer et al., 2022). Homology grouping was found to be optimal (highest F1-score) at the ‘D3’ setting based on the distribution of ‘eudicots_odb10’ (BUSCO) orthologues across homology groups. As no method existed to integrate called PAVs across genomes (*i*.*c*. proteomes), we developed novel functionality to add bPAV information to the graph database (‘add_pavs’; included in PanTools v4.2.0 and onwards). The bPAV information has to be in “binarised” format, *i*.*e*. each transcript has to be either present (‘1’) or absent (‘0’). This allows us to calculate the number of core and accessory transcripts in the super-pangenome and provide an overview of the gene space for all 474 accessions based on homology. Finally, we also extended the PanTools pangenome growth and gene classification functions to use bPAV information, from PanTools v4.2.0 onwards. Based on gene distances between the accessions, a neighbour-joining tree was generated to gain insight in the phylogeny of all four species.

### GWAS

To associate transcript presence/absence and copy numbers with phenotypes, we used the PAV and CNV matrices for *L. sativa*. Since many transcripts show no variation (Figure 2), we first reduced the size of both PAV and CNV matrices by removing those transcripts that show little to no variation in the following way: We created bins per one decimal for PAV values and we rounded CNV values to the closest integer value (these bins and rounded values are only used for the purpose of filtering out uninformative transcripts). Next, depending on availability of phenotypes for a given accession, we removed those accessions for which no phenotype value was recorded and subsequently applied two filters to the bins: 1) We removed all transcripts that showed no variation across accessions. 2) We removed all transcripts where more than 95% of all accessions had the same (rounded) value (corresponding to a minor allele frequency of 5%). Phenotypes were retrieved from the CGN database and from Mehrem et al. (2024).

GWAS was then performed with the two (filtered) matrices: PAVs and CNVs. The kinship (based on PAV) was calculated as the covariance of the genotypes, using the cov function in R v4.2. GWAS was conducted through a linear mixed model with the kinship included as random effect, using the lme4QTL package in R (Ziyatdinov et al., 2018). Bonferroni multiple testing correction was used, calling associations significant at *p <* (0.05*/*number of genotypes).

## Supporting Information

### Supplementary Methods

Additional information about the key steps we took in the methods and the reasoning behind them.

### Supplementary Data

at 4TU.ResearchData (DOI: https://www.doi.org/10.4121/c7935d6a-d6ae-42e7-af7e-0ae8cddf70d7.v1):

**SD1:** Linear pangenome sequences

**SD2:** Linear pangenome gene annotation

**SD3:** *Lactuca* AUGUSTUS gene models

**SD4:** PAV table per species

**SD5:** CNV table per species

**SD6:** Homology table across species

**SD7:** bPAV table across species

**SD8:** Full functional enrichment results core *Lactuca* genes

**SD9:** Full functional enrichment results variable *Lactuca* genes

**SD10:** Full GWAS results

**SD11:** Neighbour-joining tree with bootstrapping values

## Availability of data and materials

Additional WGS for this study have been deposited in the European Nucleotide Archive (ENA) at EMBL-EBI under accession number PRJEB63589 https://www.ebi.ac.uk/ena/browser/view/PRJEB63589. The pipeline to create the linear pangenomes is available at https://github.com/LettuceKnow/linear_pangenome_building. PanTools is available at https://git.wur.nl/bioinformatics/pantools. We provide a webportal for easy downloading, querying and visualising the data presented here: https://www.bioinformatics.nl/lettuce/.

## Authors’ contributions

**Dirk-Jan M. van Workum**: Conceptualisation, Methodology, Software, Formal Analysis, Data Curation, Writing – Original Draft, Visualisation; **Sarah L. Mehrem**: Methodology, Software, Formal Analysis, Writing – Review & Editing, Visualisation; **Basten L. Snoek**: Software, Writing – Review & Editing, Supervision; **Marrit C. Alderkamp**: Resources, Data Curation; **Dmitry Lapin**: Resources, Data Curation, Writing – Review & Editing; **Flip F.M. Mulder**: Resources, Data Curation; **Guido F.J.M. Van den Ackerveken**: Resources, Data Curation, Writing – Review & Editing, Funding acquisition; **Dick de Ridder**: Writing – Review & Editing, Supervision; **M. Eric Schranz**: Writing – Review & Editing, Supervision; **Sandra Smit**: Conceptualisation, Methodology, Writing – Review & Editing, Supervision

## Acknowledgements

We would like to thank Eef Jonkheer and Robin van Esch for their support in the implementation of novel PanTools functionalities.

## A Supplementary Figures

**Supplementary Figure S1:**
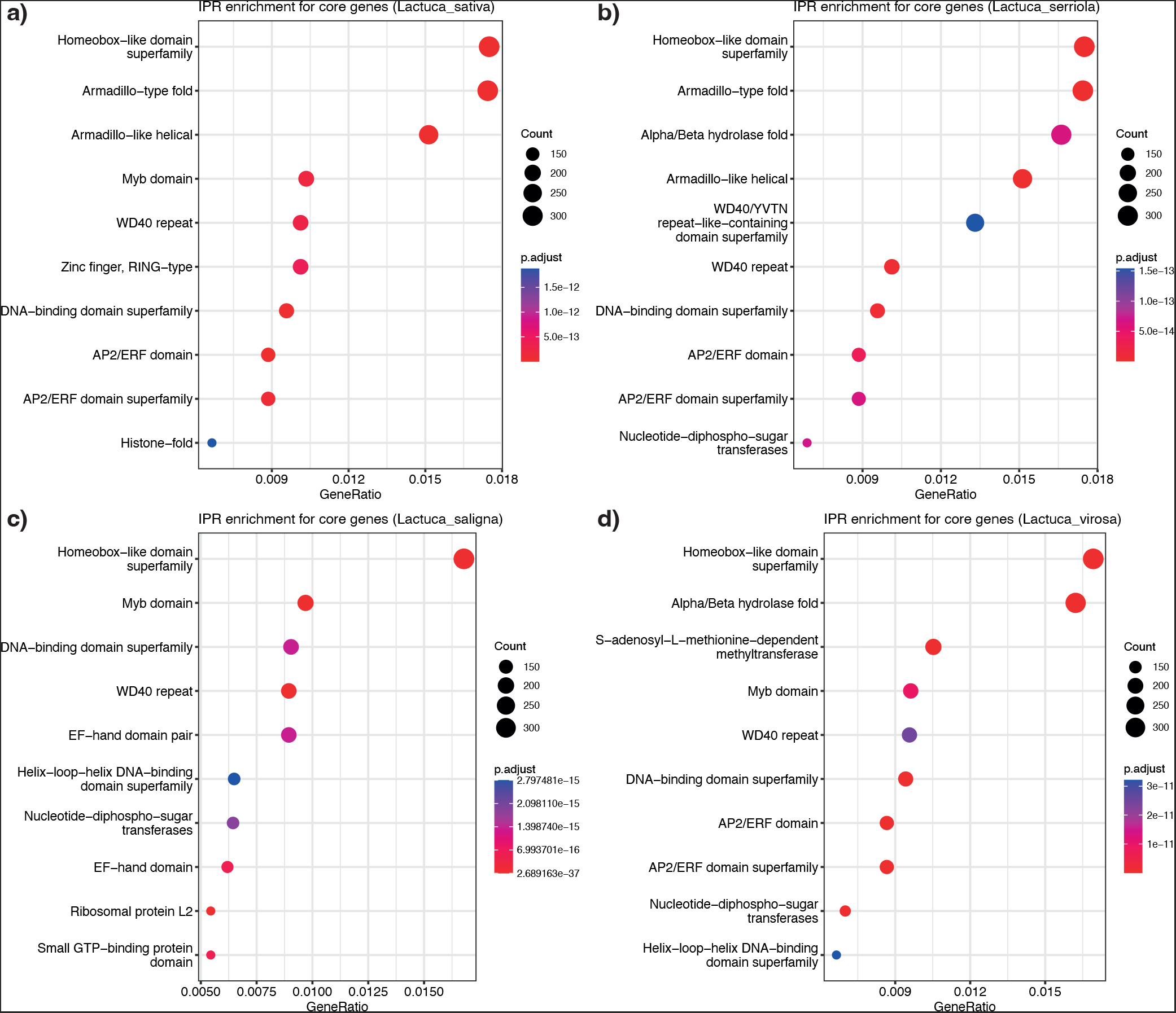
Functional enrichment of the core *Lactuca* genes in *L. sativa* (**a**), *L. serriola* (**b**), *L. saligna* (**c**) and *L. virosa* (**d**) using InterPro domains. Only the first ten most significant domains are shown (p-value adjusted according to Bonferroni). For full results, see Supplementary Data 8.

**Supplementary Figure S2:**
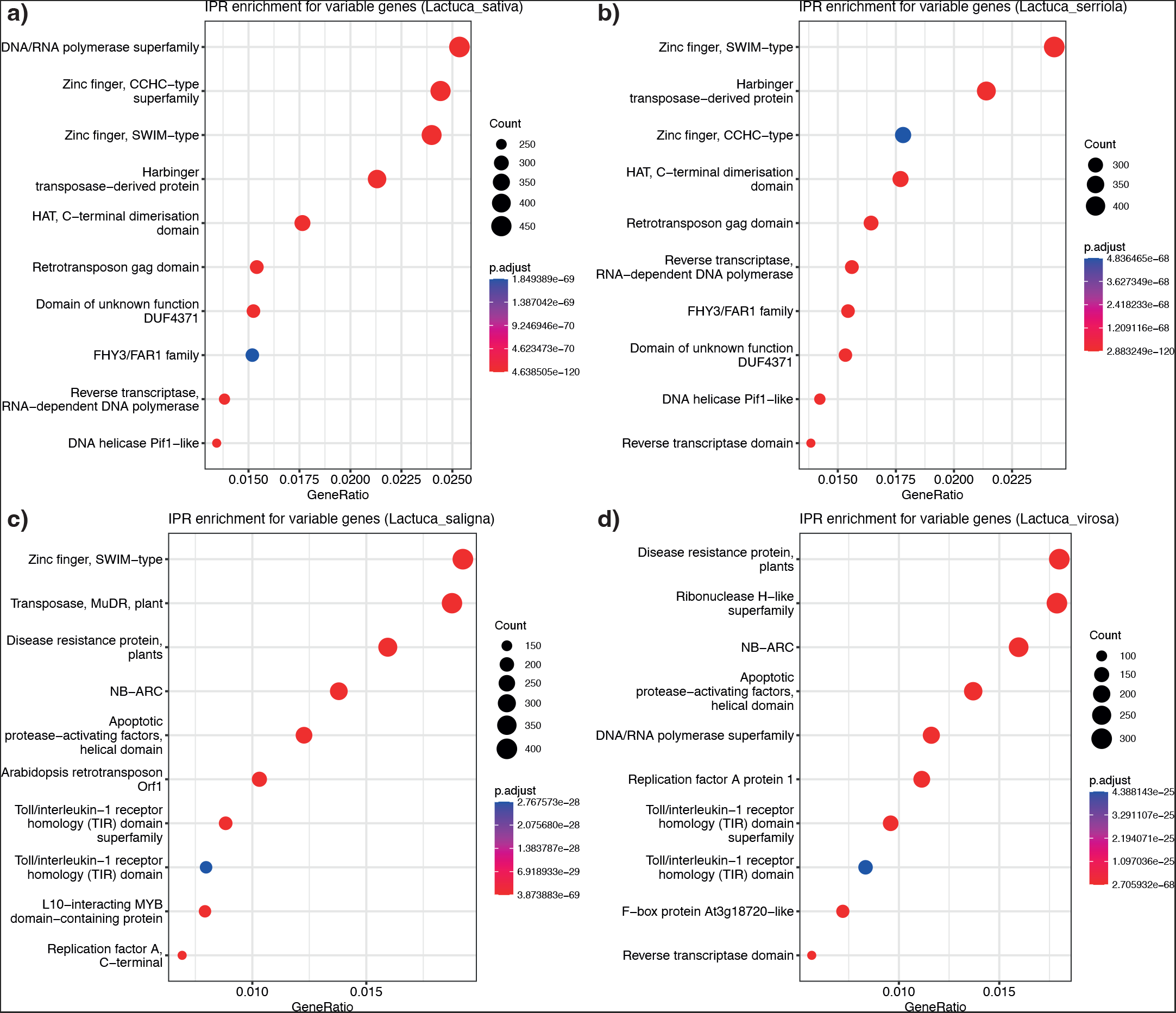
Functional enrichment of the variable *Lactuca* genes in *L. sativa* (**a**), *L. serriola* (**b**), *L. saligna* (**c**) and *L. virosa* (**d**) using InterPro domains. Only the first ten most significant domains are shown (p-value adjusted according to Bonferroni). For full results, see Supplementary Data 9.

**Supplementary Figure S3:**
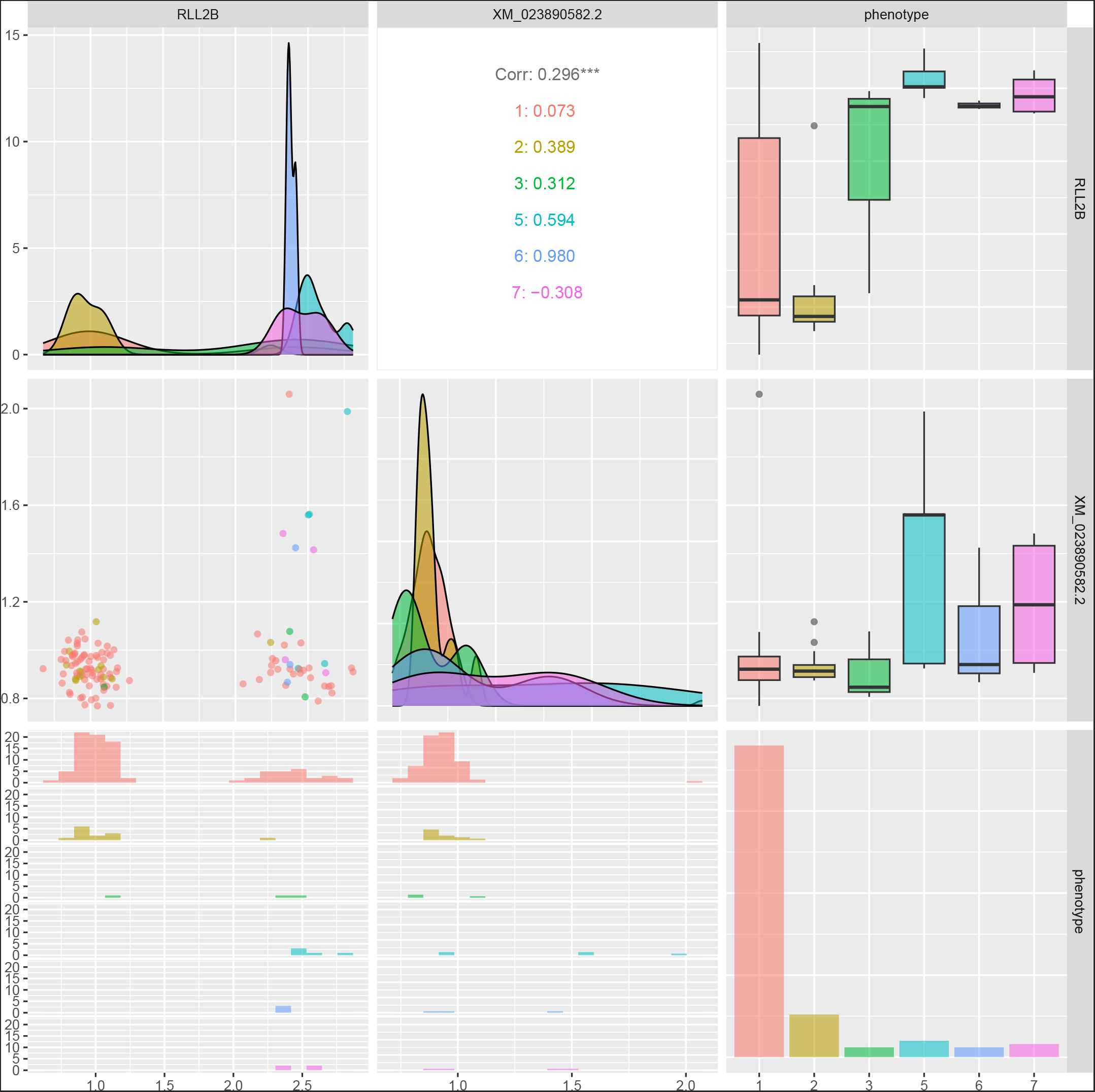
Correlation of CNV and phenotype values for the “LeafAnthocyaninContent” CNV-GWAS peak on chromosome 5 is shown here. Column/row 1 has the CNV for *RLL2B* (XM_023889304.2), column/row 2 shows CNV for the most significant mRNA hit (on chromosome 5): XM_023890582.2 and column/row 3 shows the “LeafAnthocyaninContent” phenotype from CGN. The diagonal shows the distribution of CNV and phenotype values for their respective columns/rows. Phenotype values are discrete numbers and therefore shows as barplot instead. The bottom triangle shows all correlation plots between CNV values and phenotype. From these it can be seen that the CNV-GWAS hits is correlated to the *RLL2B* gene. This corresponds to the correlation of *RLL2B* CNV values with the anthocyanin phenotype (top right), indicating that a higher copy number of *RLL2B* indeed correlates with a higher leaf anthocyanin content. This plot was created with the ‘ggpairs’ function from the R package “GGally”.

**Supplementary Figure S4:**
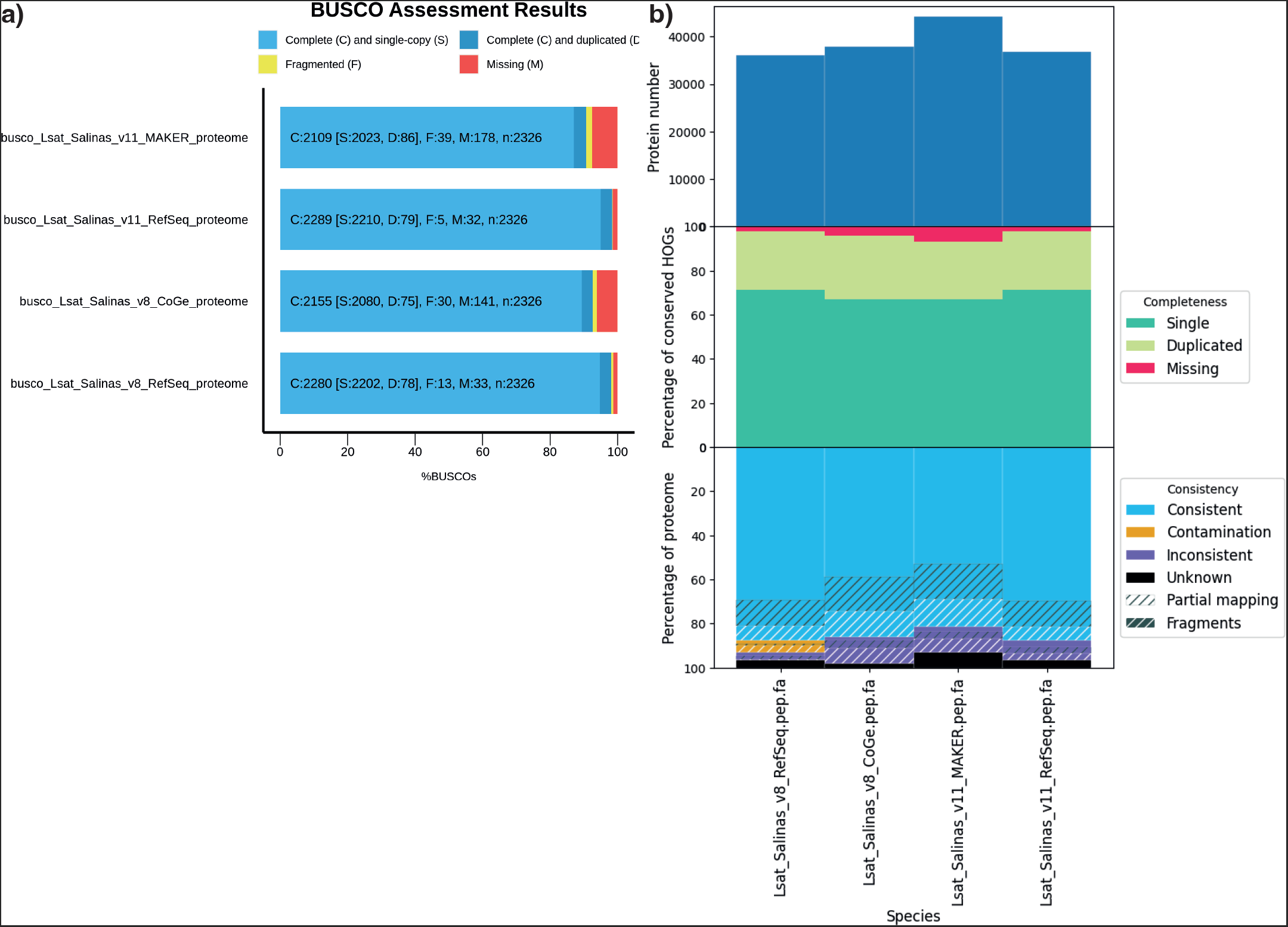
Completeness comparison of the proteome for *L. sativa* var. Salinas v8 and v11 with BUSCO v5.2.2 (**a**) and OMArk v0.2.3 (**b**) (Manni et al., 2021; Nevers et al., 2024). For both v8 and v11, the original annotation (“CoGe” and “MAKER”, respectively) and the annotation as generated by RefSeq are included.

