## Supplementary methods for "*Lactuca* super-pangenome reduces bias towards reference genes in lettuce research"

**PAV batch effect** **11**

### Constructing a linear pangenome

#### Data selection

Constructing a linear pangenome involves mapping resequenced accessions to a chosen reference assembly. At the time of starting the analyses, we had access to the following genome assemblies:

- *Lactuca sativa*
  - Salinas v8 (CoGe: 28333)
  - Salinas v8 (RefSeq: GCF_002870075.2)
  - Tizian (CoGe: 36441)
- *Lactuca serriola*
  - US96UC23 (CNSA: CNP0000335 (TKI-340))
  - US96UC23 (CoGe: 30885; personal communication with Richard Michelmore, UC Davis)
- *Lactuca saligna*
  - CGN05271 (CNSA: CNP0000335 (TKI-342))
  - CGN05327 (GenBank: PRJEB56287)
- *Lactuca virosa*
  - CGN04683 (CNSA: CNP0000335 (TKI-404))
  - CGN04683 (GenBank: GCA_928212115.1)

We compared completeness and mapping statistics for all of these assemblies and their annotations (Supplementary Methods Figure 1, Supplementary Methods Table 1). Based on these results, we decided to use the following references for read mapping: *L. sativa* Salinas v8 (GCF_002870075.2) for *L. sativa* and *L. serriola* accessions, *L. saligna* CGN05327 (PRJEB56287) for *L. saligna* accessions and *L. virosa* CGN04683 (GCA_928212115.1) for *L. virosa* accessions.

Next to these high-quality genome assemblies, we used the short-read sequencing data from Wei et al. (2021) and 68 additionally sequenced accessions (Supplementary Table B).

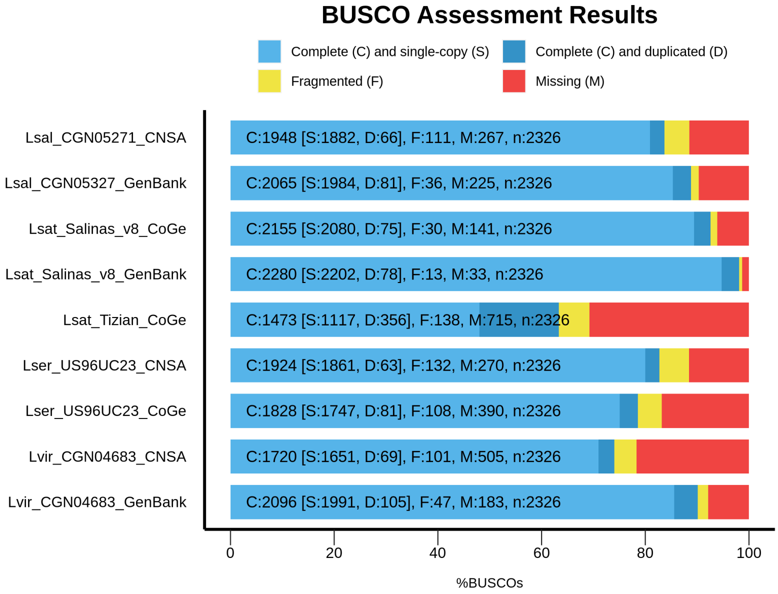

Supplementary Methods Figure 1: BUSCO completeness assessment for available *Lactuca* proteomes in alphabetical order. “Lsat” means *L. sativa*, “Lser” *L. serriola*, “Lsal” *L. saligna*, and “Lvir” *L. virosa*.

As noted already by Wei et al. (2021) in their Supplementary Table 5, some accessions are likely crosses between multiple species or wrongly labelled. Based on their findings, we sequenced LK450 (PV15242, 275-5) again for *L. saligna* and we removed LK467 from the set for *L. virosa* as this appeared to be a *Lactuca georgica*. Furthermore, we redid the analysis Wei et al. (2021) did for *L. sativa* and *L. serriola* for discerning which accession belongs to which species. We created a covariance matrix of the previously identified single-nucleotide

Supplementary Methods Table 1: Pair-wise mapping statistics for the most complete genome assemblies per species to detect any outliers. For *L. sativa* we used GCF_002870075.2 (RefSeq), for *L. serriola* 30885 (CoGe), for *L. saligna* PRJEB56287 (GenBank), and for *L. virosa* GCA_928212115.1 (GenBank). For each set of WGS data, we randomly selected one pair of fastq files for each reference accession and mapped it to the reference genome using bwa-mem2 (after trimming with Trimmomatic; *cfr.* Methods). The highest number per row is highlighted, indicating the best genome assembly for this WGS data.

| | Reads mapped to genome assembly (properly paired) | | | |
|---|---|---|---|---|
| WGS data | *L. sativa* | *L. serriola* | *L. saligna* | *L. virosa* |
| CNS0047520 (*L. sativa*) | **99.8%** (**97.1%**) | 98.6% (79.7%) | 95.6% (73.4%) | 94.3% (75.0%) |
| CNS0047707 (*L. serriola*) | **99.4%** (**92.1%**) | 99.0% (80.5%) | 95.7% (71.9%) | 94.7% (74.2%) |
| CNS0047873 (*L. saligna*) | 96.4% (76.4%) | 95.6% (67.0%) | **99.1%** (**95.5%**) | 92.4% (71.3%) |
| CNS0047910 (*L. virosa*) | 94.8% (67.7%) | 94.3% (65.4%) | 93.0% (62.7%) | **98.0%** (**93.2%**) |

polymorphisms (SNPs) (Supplementary Methods Figure 2) and a covariance matrix of the presence/absence variation (PAV) identified in this study (Supplementary Methods Figure 3). From these matrices, we identified 9 accessions that are either crosses or mis-labelled (Supplementary Table B). Thus, we end up with a total of 474 accessions, including 406 accessions from Wei et al. (2021) and 68 accessions that we sequenced as part of the LettuceKnow project.

#### Pipeline construction strategy

For constructing a *Lactuca* super-pangenome, we considered several approaches:

- constructing a graph-based pangenome from 474 *de novo* assembled lettuce accessions
- constructing one linear pangenome for all 474 lettuce accessions
- constructing one linear pangenome per species and combining these in a graph super-pangenome

Since the available resequencing data was sequenced with Illumina (short-read) at a depth of about 20X per accession and since lettuce has a large genome size, we reasoned that *de novo* assembly will not give us good quality genomes for pangenomics. The fragmented *de novo* assemblies from Wei et al. (2021), which were sequenced at a higher coverage, confirm this. Since the availability of high-quality assemblies is essential for pangenomics, we chose not to follow this first approach. Next, creating one linear pangenome for all accessions (belonging to four species) will make pangenomics difficult as well because of the large diversity between the species: *L. sativa*, *L. saligna* and *L. virosa* only share about half of their genes (Xiong et al., 2023). Therefore, we decided to build on the idea of a super-pangenome as proposed by Khan et al. (2020) and already implemented for *e.g.* rice (Shang et al., 2022). We implemented this by first constructing linear pangenomes for each of the four species based on resequencing data and then integrating the four linear pangenomes in a graph super-pangenome based on PAV.

Even though most papers describing a linear pangenome use a custom pipeline, so far there have been two efforts (to our knowledge) for the standardisation of eukaryotic pangenome creation pipelines (Glick & Mayrose, 2021; Hu et al., 2017). However, these pipelines seem to have been developed for species with a smaller genome size than lettuce. Typical showcases are yeast and *Arabidopsis thaliana*, both at least 20 times smaller than lettuce. Keeping this in mind, we tested the newer Panoramic but found out that it could not handle the large size of lettuce in reasonable time and that it did not allow for data-specific contamination filtering steps (Glick & Mayrose, 2021). Therefore, we decided to create a custom pipeline for the creation of linear pangenomes. We implemented this pipeline in Snakemake, building on the map-to-pan idea of earlier published approaches (Glick & Mayrose, 2021; Golicz, Batley, et al., 2016). This pipeline can be found under github.com/LettuceKnow/linear_pangenome_building.

The purpose of our lettuce super-pangenome is the creation of an accurate overview of all genic variation in both cultivated and wild lettuce. Therefore, we heavily rely on an accurate gene annotation in our analyses. Since most graph (pan)genome tools create a graph in GFA (or similar) format which is difficult to analyse gene annotations in, we decided to use PanTools for our analyses (Sheikhizadeh et al., 2016). PanTools stores all information in an interactive neo4j graph database. Moreover, next to gene annotations, PanTools allows the functional annotation of these genes.

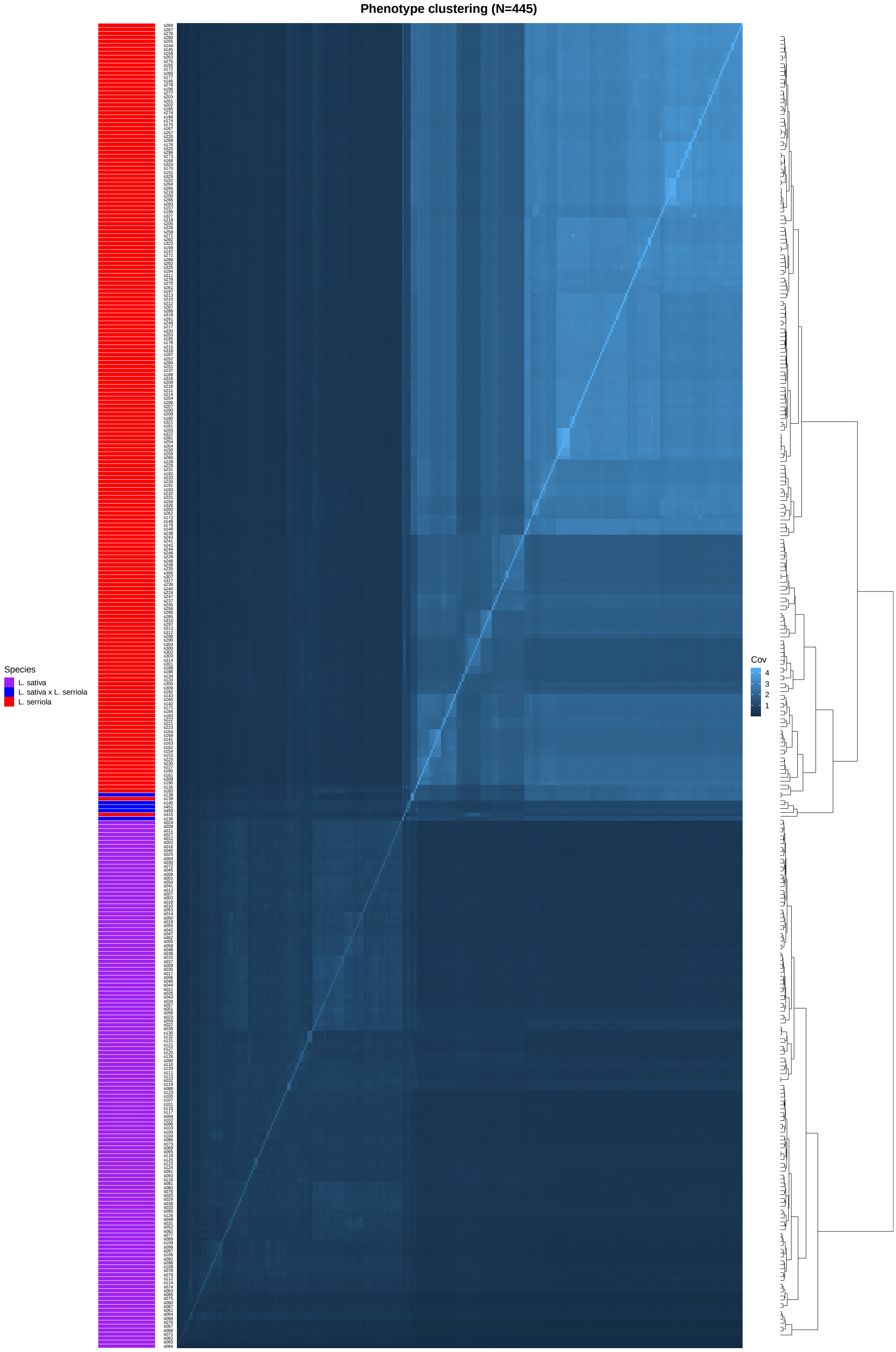

Supplementary Methods Figure 2: Kinship for all *L. sativa* and *L. serriola* accessions based on covariance of SNPs as called by Wei et al. (2021). Identifiers used here are the original identifiers, which can be converted to other identifiers (Supplementary Table B). This figure was used together with Supplementary Methods Figure 3 used for outlier detection.

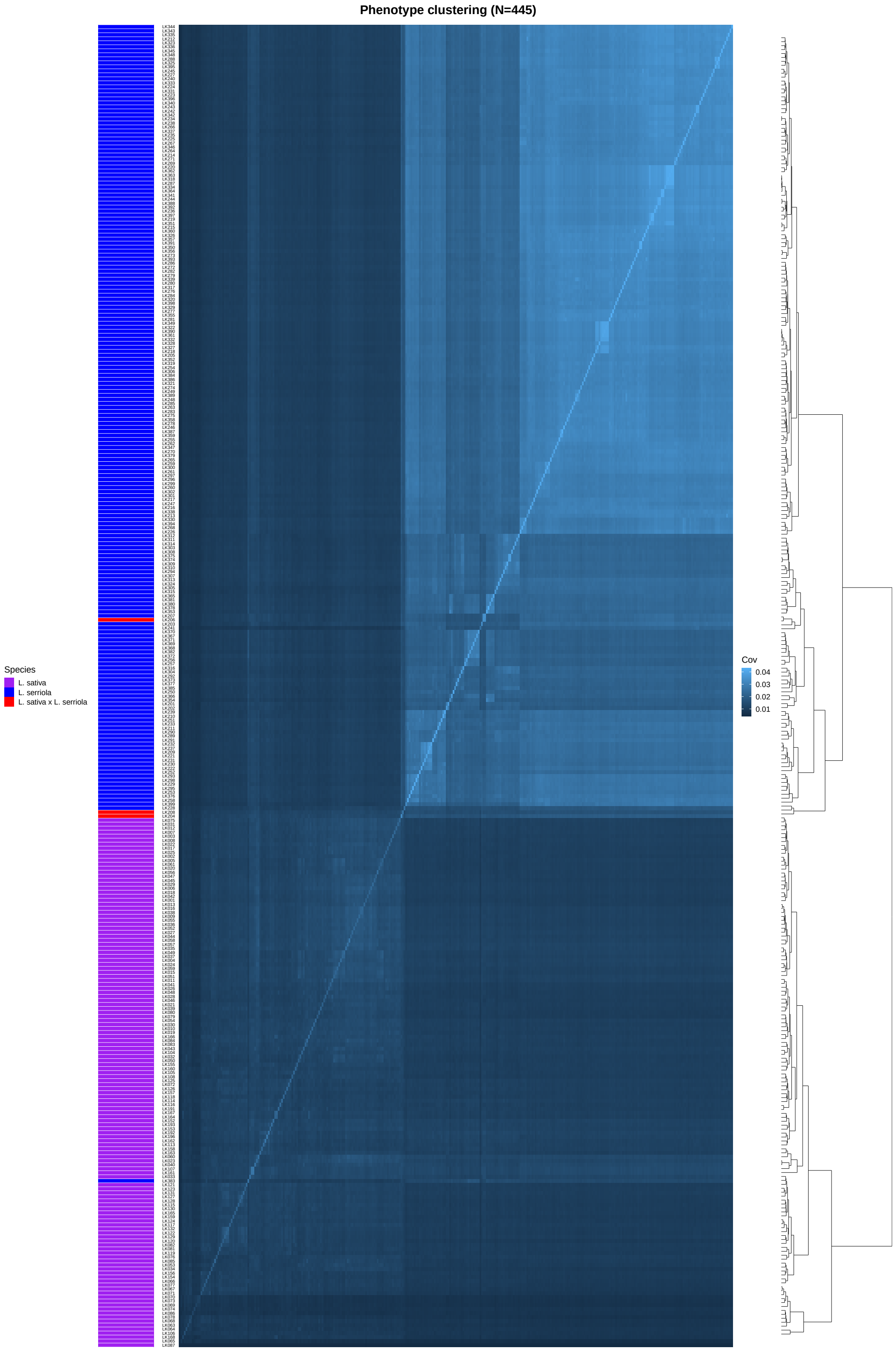

Supplementary Methods Figure 3: Kinship for all *L. sativa* and *L. serriola* accessions based on covariance of PAV values. This figure was used together with Supplementary Methods Figure 2 used for outlier detection.

#### Linear pangenome construction strategy

For the construction of linear pangenomes, roughly two strategies exist (Bayer et al., 2020; Golicz, Batley, et al., 2016): *de novo* assembly and iterative assembly. As reasoned above, *de novo* assembly is not possible with the available data. Therefore, we tested two variations dealing with only unmapped reads:

- a "map-and-iteratively-assemble" approach, where we iteratively add assembled unmapped reads after mapping all libraries in parallel to the same reference genome.
- a "co-assembly"-like approach, where we assemble all sets of unmapped reads (after mapping all libraries in parallel to the same reference genome) into one assembly.

We did not test an "iterative-map-and-assemble" approach since the mapping of libraries cannot be done in parallel here, taking too much time for the size of our data.

For testing purposes, we combined the complete set of *L. sativa* and *L. serriola* accessions and used GCF_002870075.2 (*L. sativa*) as a reference genome. The resulting novel sequence for the two approaches tested, differed by roughly a factor 2, both in terms of sequence length and number of annotated genes (the "co-assembly"-like approach being the larger one). Interestingly, when mapping the input reads for the assembly process back to the assemblies (for four random accessions), we found no difference (Supplementary Table 2). We therefore investigated the difference in assembly size by performing several filtering steps. Filtering genes on contamination or sequences on length or GC-content all had no difference on the factor 2 difference between the two strategies (Supplementary Table 2). Only a small difference in the GC-content distribution could be identified (Supplementary Methods Figure 4). Thus, since the "map-and-iteratively-assembly" approach was just as complete (in terms of mapped reads) but smaller in size, we chose this strategy.

#### Contamination filtering

When performing short-read sequencing on plant tissues, it is next to impossible to get uncontaminated libraries. Since one essentially sequences the microbial community of the plant tissues as well, we reasoned that the problem of separating contamination from actual novel *Lactuca* sequence is similar to the binning problem in metagenomics. A popular tool used in metagenomics for the classification of contigs is kraken2. We believe this tool can be applied to the assembly of unmapped reads per accession as well to filter out large taxonomic groups completely unrelated to *Lactuca*. Since kraken2 is fast (and typically has a large amount of no-hits), we used kraken2 for the classification of the newly produced assembly of every set of unmapped reads (thus, per accession) against the NCBI nt database. Testing this for the linear pangenome of *L. sativa* and *L. serriola*, the largest taxonomic groups identified in this contamination check were bacteria, some (pest) insects such as thrips and whitefly, and amoebae.

Since up to 6.5% of sequences has no kraken2 hit and a large part of the hits were bacteria, we calculated GC content per sequence to see whether we could remove large parts of the bacterial content based on GC content. First, we determined the GC content of *Lactuca* by dividing the *L. sativa* genome in parts of 1kbp. The average GC content of the *L. sativa* reference genome is 34% (a maximum of about 50% and clearly unimodal) whereas the GC content of the novel sequences is bimodal with an additional large peak just above 50%

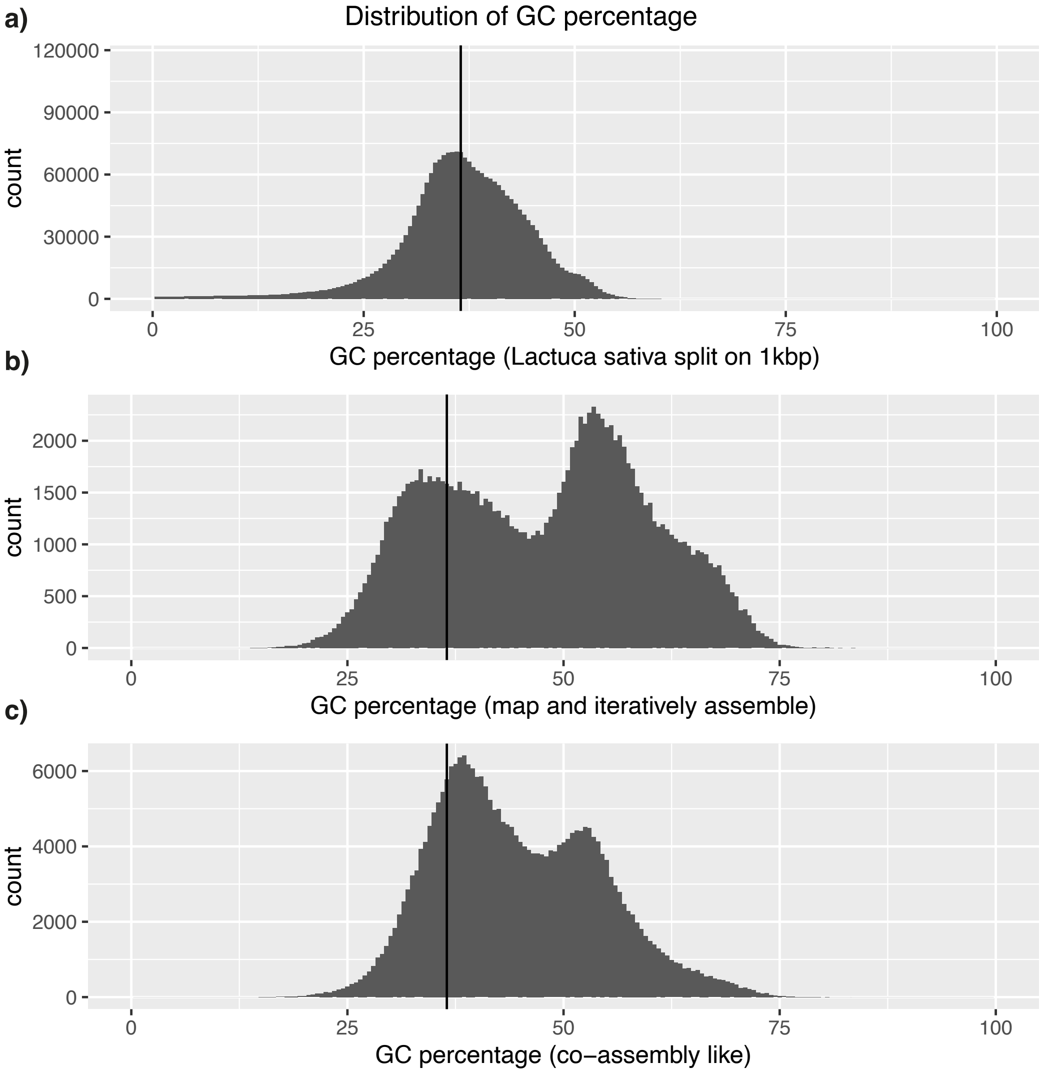

Supplementary Methods Figure 4: GC distribution for 1kbp sequences in the *L. sativa* reference genome (**a**), the novel sequence content after the map-and-iteratively-assemble approach (**b**) and after the co-assembly-like approach (**c**). The vertical black line indicates the median value for *L. sativa* (36.5%).

(Supplementary Methods Figure 4). Thus, we decided to remove all sequences with a GC content above 50%.

After the iterative process of combining all the assemblies, we created a blobplot for the combined set of novel sequences using blobtools v1.1.1 (Laetsch & Blaxter, 2017). Even though filtering based on GC content and a kraken2 search was already performed, still non-plant sequences could be identified with a blastn search against nt. Most of the contamination left in the pangenome was other eukaryotes (Supplementary Methods Figure 5). Based on the blobplot, we performed a final filtering step in which we filtered out all sequences that have a blastn hit that was neither “Streptophyta” or “no-hit”. Also, we reasoned that all sequences with a coverage lower than 10X could safely be removed without deleting real lettuce sequences: the coverage was calculated based on all accessions that separately had an average coverage of about 20X.

Finally, we performed one additional search for contamination: with the predicted genes. It is easier to find hits for protein sequences since those are typically more conserved than nucleotide sequences. Therefore, we did a final search of the novel proteins against the nr database with mmseqs2. From the 7,843 predicted genes in our test dataset with *L. sativa* and *L. serriola*, we found that 5,766 were either unclassified or classified as eukaryotic. We kept all sequences that either had no genes annotated or had at least one eukaryotic or unclassified gene.

#### Duplication filtering

Aiming for a non-redundant but complete linear pangenome, it is essential to filter well for duplications. Various strategies have been used in the past for filtering out redundancy in linear pangenome construction. In the *Brassica* pangenome of Golicz, Bayer, et al. (2016), an “iterative-map-and-assembly” approach was used. For each iteration, the newly assembled contigs were aligned to the reference genome with “LASTZ (–notransition –chain –ambiguous=n –identity=93 –coverage=90 –continuity=95)” to prevent any contig to be added more than once. Gao et al. (2019) built their tomato pangenome with a similarity cutoff of 90% at every step for removing redundancy. This threshold is based on an analysis they did where they plotted the cutoff against the percentage non-reference sequence in their pangenome. They found that the percentage non-reference sequence was stable up to 80%, increased linearly between 80% and 90% and increased exponentially above 90%. Finally, Hübner et al. (2019) used two different thresholds in the sunflower pangenome: they 1) filtered out all novel sequences that have >75% similarity over >75% length when aligned against the reference genome 2) removed redundancy from the set of novel sequences by applying a 95% cutoff. We used these existing approaches as inspiration for this step in our pipeline.

We filter out redundancy at two stages: during the iterative building process and during a final clustering of the novel sequences. We checked that the assembled novel sequences are truly absent from the reference genome by including the entire reference genome in the iterative assembly process. In the test dataset of *L. sativa* and *L. serriola* with GCF_002870075.2 (*L. sativa*) as a reference genome, we found no hits of the novel sequences against the reference sequences but the total number of novel sequences was reduced from 403,770 to 17,417. Next, we applied a 90% clustering cutoff on the novel sequences which reduced the number of novel sequences to 17,014. We reasoned that if there were large redundancy in our pan-

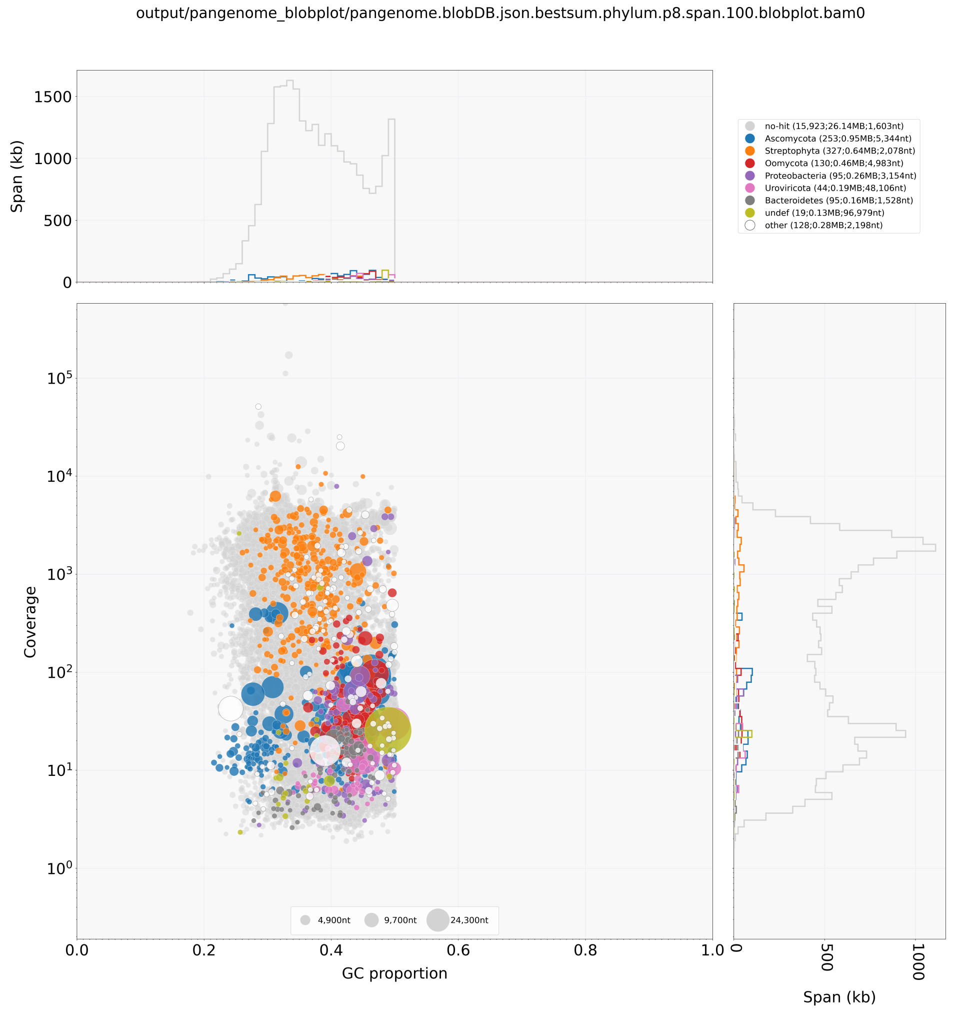

Supplementary Methods Figure 5: Blobplot for all novel sequences in a test dataset of *L. sativa* and *L. serriola* for determining the final contamination filtering step. The coverage is determined based on the mapping of the unmapped reads of all libraries combined and the taxonomy is based on the best blastn hit against the nt database.

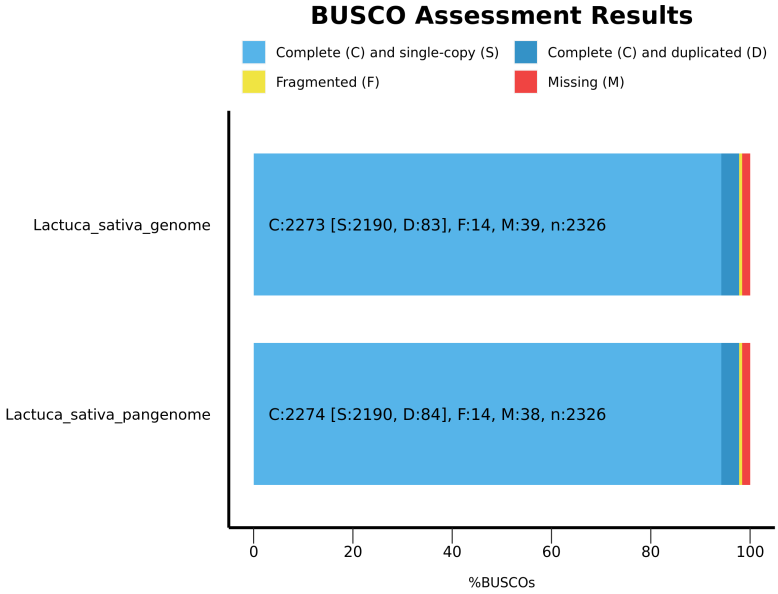

Supplementary Methods Figure 6: BUSCO scores for the reference genome of *L. sativa* and the pangenome created from the test dataset (both *L. sativa* and *L. serriola* accessions) for determining possible duplications in the final pangenome.

genome, we should be able to see this in an increased number of duplicated BUSCO genes. This was however not the case (Supplementary Methods Figure 6).

#### Gene prediction

As we did not have access to RNA-seq data for multiple tissues of all accessions, it was difficult to accurately annotate novel genes in the novel sequence content. Therefore, we only did simple gene prediction and kept in mind that genes showing potentially interesting biology first need to be validated experimentally (as we would recommend for reference genes too). For gene prediction, we tested three different AUGUSTUS models: 1) the *A. thaliana* model, 2) the tomato model, and 3) a custom made *Lactuca* model. For this last model, we used the set of conserved genes between *L. sativa*, *L. saligna* and *L. virosa* as input for training AUGUSTUS (using 1kbp of sequence up- and downstream of the genes, 100 genes for testing, and the other genes for training). We found that the custom *Lactuca* model slightly outperformed the *A. thaliana* and tomato models and therefore decided to use our *Lactuca* model for gene prediction (Supplementary Methods Table 3).

### PAVs and CNVs

#### PAV and CNV calculation

We calculated PAV values based on the average **coverage** (horizontal) of the exon regions of a transcript and copy-number variation (CNV) values based on the average (normalised) **depth** (vertical) of the exon regions of a transcript. Here, we defined transcripts as any genomic feature annotated in the gff3 file with exons. Initially, we planned on discerning absent genes, present genes and multi-copy genes all based on the depth. Since the average depth of all transcripts of one accession depends (mostly) on the average genome coverage, comparing between accessions is impossible since the average genome coverage differs. Therefore, we normalised the depth values by dividing each value by the median of all depth values per accession. Nevertheless, a cutoff between absent and present genes based on depth appeared to be difficult to set with our data, let alone setting a cutoff between single- and multi-copy genes (*e.g.* LK198; Supplementary Methods Figure 7a). Discerning between absence and presence based on coverage gave more promising results (even though we were now unable to discern between single- and multi-copy genes) (*e.g.* LK198; Supplementary Methods Figure 7b). Therefore, our analyses were mostly based on coverage (*i.e.* PAV). Although absence, presence and copy-number variations may be separable on the level of individual genes, we only use depth for CNV.

Setting a threshold for absence/presence based on the underlying biology or the distribution of PAV values for genes appeared to be difficult. Others have used a wide range of values (see Discussion). Since it is not only visually difficult to set this threshold but also biologically impossible to determine whether a gene is present based on its coverage, we decided to keep the continuous values for *e.g.* genome-wide association study (GWAS). When one wants to be sure whether a specific gene is truly present or absent in any accession, one would need to look at what part of the gene is absent, whether it might have duplications and RNA-seq evidence. Even then, a partially present gene can still be transcribed, translated into protein and even have a similar function if it misses a large part of its sequence. Ultimately, the effect of a PAV can only be discovered experimentally by molecular biology.

However, some applications require a threshold to be chosen, *e.g.* pangenome growth curves and integrating PAVs across species. We therefore compared several thresholds and their effect on the total number of variable transcripts for creating a binary PAV (bPAV) matrix (Supplementary Methods Figure 8). A transcript is defined as “variable” if it is absent in at least one accession according to the threshold. We specifically considered the thresholds 0.5 as midpoint between 0 and 1, and 0.8 based on the increase of variable genes (Supplementary Methods Figure 9). 0.5 seemed to separate a linear increase and an exponential increase in the number of variable genes. Any threshold above 0.8 had a visibly large difference with “neighbouring” threshold values. We also tested the effect of both thresholds on the integrated bPAVs across species by generating a neighbour-joining tree using both and found that there was little difference between 0.5 and 0.8. Since the impact of the exact threshold was only minor, we decided to go with 0.8 because a transcript that is covered between 50% and 80% is biologically speaking more likely to be absent than present.

### PAV batch effect

When visualising the PAVs for *L. sativa*, there appeared to be a batch effect in the PAV data based on the origin of sequencing (Supplementary Methods Figure 10). Specifically, non-reference genes introduced by the LettuceKnow sequencing effort accessions seem to contain genes specific to that sequencing effort. Accessions LK099, LK014, LK088, LK141 and LK139 seem to be most affected by this. As these accessions all belong to different subgroups, it is unlikely that this is a subgroup specific set of genes. Also, no clear effect is seen in the PAV values of the reference genes in these accessions. Since we already filtered for contamination

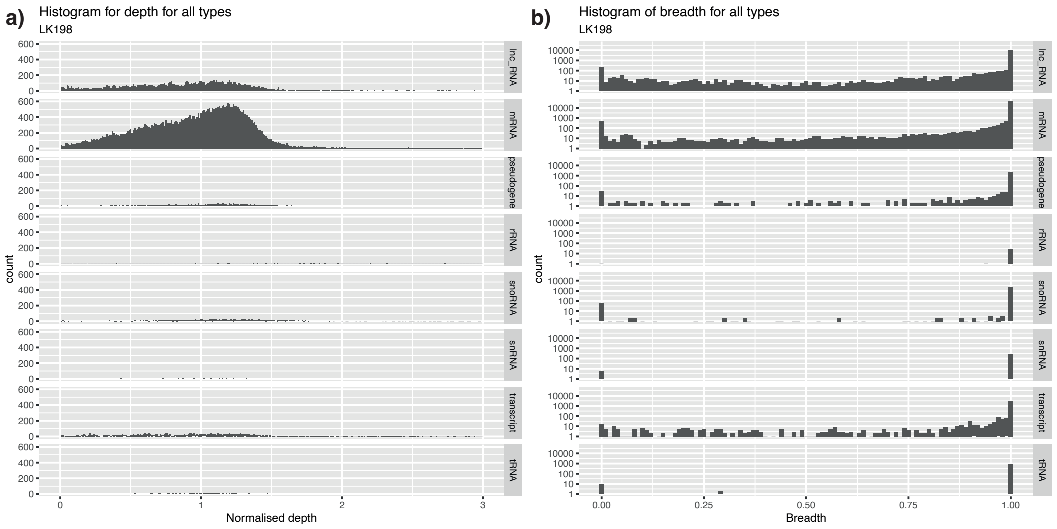

Supplementary Methods Figure 7: Distribution of depth (vertical; CNV) and breadth (horizontal; PAV) in LK198 (as an example) for all types of transcripts annotated in the *L. sativa* genome that have exons. Only exons are considered for these calculations. **a)** Depth is calculated as the average depth of a transcript and normalised by dividing by the median depth of all transcripts. **b)** Breadth is calculated as the fraction of the length of a transcript that is covered by at least one read.

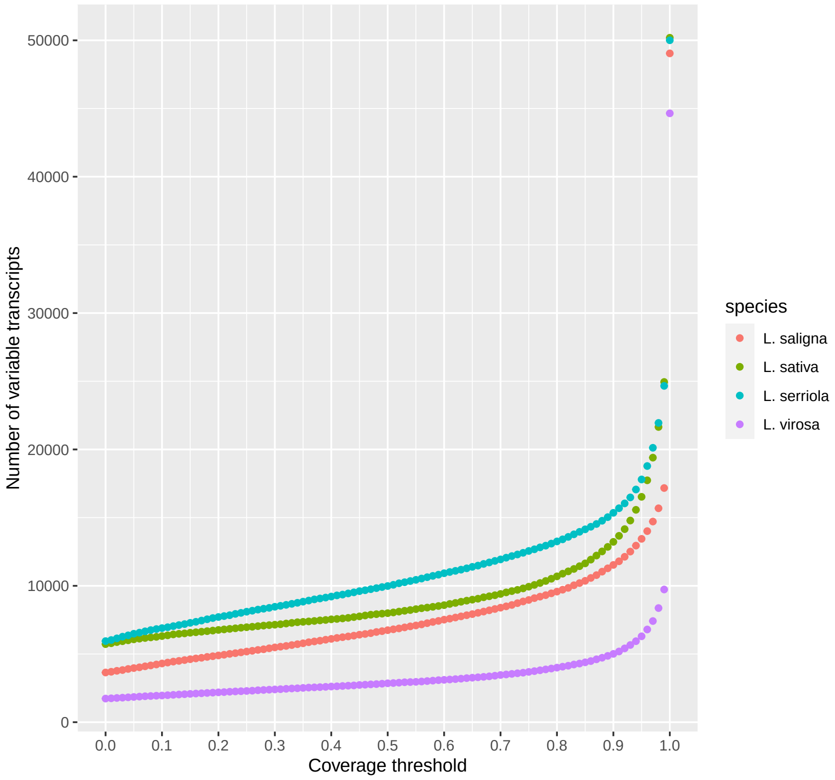

Supplementary Methods Figure 8: The effect of different coverage thresholds. For every value from 0 to 1 with a 0.01 increase, the number of variable genes is calculated per species. A variable gene is a gene that is absent in at least one accession according to the threshold.

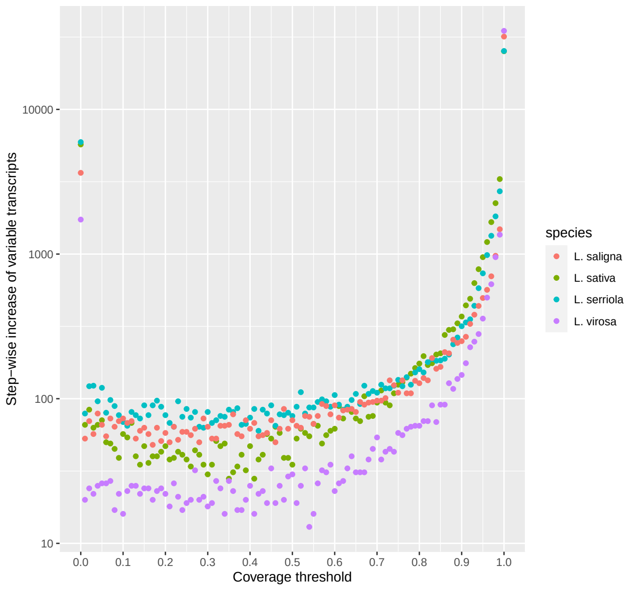

Supplementary Methods Figure 9: The step-wise increase of the effect of different coverage thresholds. For every value from 0 to 1 with a 0.01 increase, the number of variable genes is calculated per species (*cfr.* Supplementary Methods Figure 8). A variable gene is a gene that is absent in at least one accession according to the threshold. The increase is calculated as the difference in the number of variable genes with the threshold 0.01 lower.

with every known database, it is hard to distinguish whether the genes causing this batch effect are true contamination or actual lettuce genes. A run with Whokaryote (Pronk & Medema, 2022) and Tiara (Karlicki et al., 2022) suggests that they are eukaryotic genes, however. Thus, we did not filter out these genes but treated these affected accessions with care.

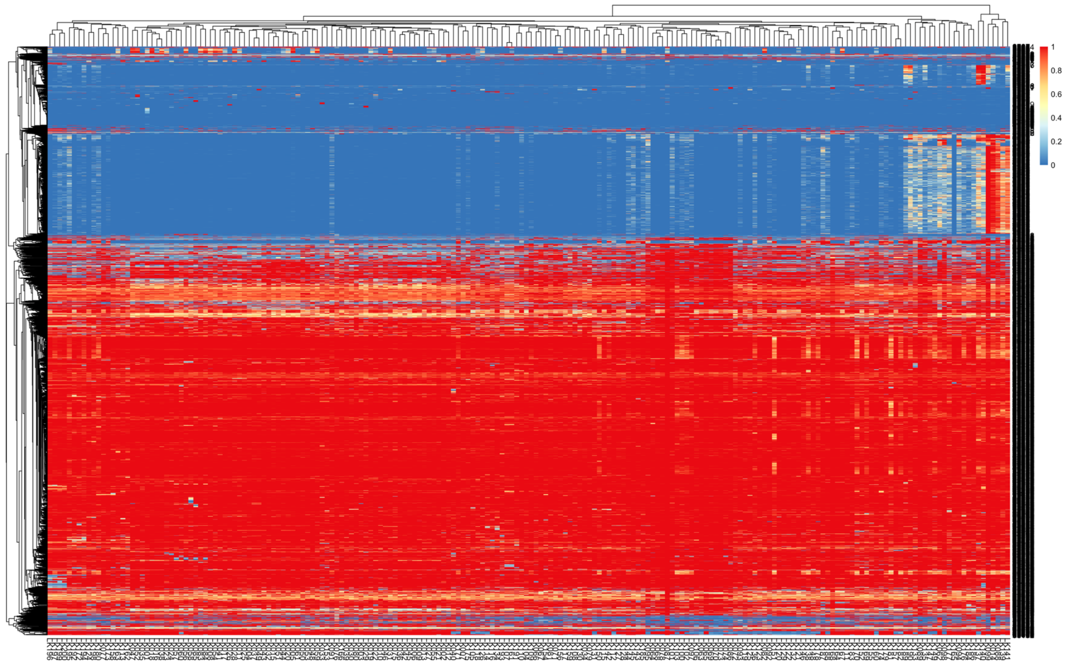

Supplementary Methods Figure 10: Heatmap of PAV values of all variable genes (rows) for all *L. sativa* accessions (columns). Variable genes here are defined as genes for which the PAV value is below 0.8 in at least one accession.

Supplementary Methods Table 2: Assembly, annotation and mapping statistics for a comparison of the “Map and iteratively assembly” versus the “Co-assembly like” approach of constructing a linear pangenome. All statistics are calculated for the novel part in the linear pangenome (*i.e.* those sequences added by the approach).

| Metric | Map and iteratively assemble | Co-assembly like |
|---|---|---|
| Total length (bp) | 133,056,926 | 272,837,691 |
| #Sequences | 125,407 | 272,172 |
| Min length (bp) | 500 | 500 |
| Avg length (bp) | 1,061 | 1,002.4 |
| Max length (bp) | 329,111 | 328,306 |
| masked | 8.98% | 6.62% |
| #Genes | 110,686 | 193,616 |
| #Genes (complete) | 28,783 | 56,168 |
| #Genes (len(seq) > 1kb) | 42,49 | 80,227 |
| #Genes (len(seq) > 5kb) | 11,237 | 19,00 |
| #Genes (GC(seq) < 50%) | 29,408 | 81,182 |
| #Genes (GC(seq) < 50% AND len(seq) > 1kb) | 10,370 | 27,549 |
| #Genes (GC(seq) < 50% AND len(seq) > 5kb) | 1,101 | 2,33 |
| LK001 unmapped reads | | |
| Error rate | 1.72% | 1.86% |
| Mapped | 5.1% | 5.2% |
| Proper pairs | 4.8% | 4.7% |
| MapQ 0 reads | 0.1% | 0.2% |
| Total seqs (million) | 9.7 | 9.7 |
| LK101 unmapped reads | | |
| Error rate | 0.30% | 2.36% |
| Mapped | 36.9% | 2.9% |
| Proper pairs | 30.2% | 2.4% |
| MapQ 0 reads | 0.2% | 0.1% |
| Total seqs (million) | 10.3 | 10.3 |
| LK201 unmapped reads | | |
| Error rate | 1.68% | 1.86% |
| Mapped | 72.4% | 63.8% |
| Proper pairs | 69.7% | 60.6% |
| MapQ 0 reads | 0.7% | 0.6% |
| Total seqs (million) | 1.3 | 1.3 |
| LK301 unmapped reads | | |
| Error rate | 1.70% | 1.98% |
| Mapped | 69.6% | 62.3% |
| Proper pairs | 66.0% | 57.6% |
| MapQ 0 reads | 1.0% | 1.1% |
| Total seqs (million) | 1.8 | 1.8 |

Supplementary Methods Table 3: Evaluation in terms of sensitivity and specificity values for three different AUGUSTUS models (default *A. thaliana*, default tomato and custom *Lactuca* models) on *Lactuca* genomes.

| | *A. thaliana* | | tomato | | *Lactuca* | |
|---|---|---|---|---|---|---|
| Level | sensitivity | specificity | sensitivity | specificity | sensitivity | specificity |
| nucleotide | 0.948 | 0.953 | 0.950 | 0.951 | 0.980 | 0.929 |
| exon | 0.858 | 0.875 | 0.830 | 0.864 | 0.894 | 0.863 |
| gene | 0.510 | 0.459 | 0.420 | 0.385 | 0.540 | 0.450 |
